# Immune disease risk variants regulate gene expression dynamics during CD4+ T cell activation

**DOI:** 10.1101/2021.12.06.470953

**Authors:** Blagoje Soskic, Eddie Cano-Gamez, Deborah J. Smyth, Kirsty Ambridge, Ziying Ke, Lara Bossini-Castillo, Joanna Kaplanis, Lucia Ramirez-Navarro, Nikolina Nakic, Jorge Esparza-Gordillo, Wendy Rowan, David Wille, David F. Tough, Paola G. Bronson, Gosia Trynka

**Affiliations:** Wellcome Sanger Institute, Wellcome Genome Campus, Cambridge, UK; Open Targets, Wellcome Genome Campus, Hinxton, UK; GSK, R&D, Stevenage, UK; R&D Translational Biology, Biogen, Cambridge, MA, USA

## Abstract

During activation, T cells undergo extensive changes in gene expression which shape the properties of cells to exert their effector function. Therefore, understanding the genetic regulation of gene expression during T cell activation provides essential insights into how genetic variants influence the response to infections and immune diseases. We generated a single-cell map of expression quantitative trait loci (eQTL) across a T cell activation time-course. We profiled 655,349 CD4^+^ naive and memory T cells, capturing transcriptional states of unstimulated cells and three time points of cell activation in 119 healthy individuals. We identified 38 cell clusters, including stable clusters such as central and effector memory T cells and transient clusters that were only present at individual time points of activation, such as interferon-responding cells. We mapped eQTLs using a T cell activation trajectory and identified 6,407 eQTL genes, of which a third (2,265 genes) were dynamically regulated during T cell activation. We integrated this information with GWAS variants for immune-mediated diseases and observed 127 colocalizations, with significant enrichment in dynamic eQTLs. Immune disease loci colocalized with genes that are involved in the regulation of T cell activation, and genes with similar functions tended to be perturbed in the same direction by disease risk alleles. Our results emphasize the importance of mapping context-specific gene expression regulation, provide insights into the mechanisms of genetic susceptibility of immune diseases, and help prioritize new therapeutic targets.

## Introduction

Translating variants from genome-wide association studies (GWAS) to function provides insights into disease biology and improves treatment options ^1^. However, linking disease variants to effector genes and elucidating the mechanisms by which they affect disease is challenging, as most GWAS variants reside in non-coding regions of the genome. Disease-associated variants are enriched within enhancer and promoter regions ^2,3^, and therefore are likely to act via gene expression regulation. Colocalization analysis can determine whether GWAS loci share a common causal variant with nearby expression quantitative trait loci (eQTL), implying that gene expression changes and disease risk are both driven by the same causal variants ^4^. Thus, maps of gene expression regulation across tissues and cell types provide important resources for linking GWAS signals to effector genes. However, the interpretability of results from these resources is limited because the majority of available eQTL maps were generated from bulk tissues which include a mixture of cell types. Gene expression regulation can be cell-type specific ^5^ and transcriptional profiling of tissues at a single-cell resolution has uncovered extensive cellular heterogeneity which is masked with bulk RNA sequencing ^6^. Additionally, the available eQTL maps fail to capture dynamic gene expression regulation, which can be context specific, e.g. manifesting throughout developmental stages ^7,8^ or in response to an external stimulus ^9,10^. Therefore, mapping dynamic gene expression regulation in a disease relevant tissue at a single cell level could provide a greater resolution into the molecular mechanisms which underlie GWAS associations and drive disease risk.

We and others have previously demonstrated that variants associated with complex immune diseases are enriched in enhancers and promoters upregulated upon CD4^+^ T cell activation ^11,12^. CD4^+^ T cells undergo activation in response to antigens arising from pathogens (such as from bacterial, viral or fungal infections) or self antigens (in autoimmune reaction). CD4^+^ T cells can be broadly divided into naive cells, which have not encountered an antigen, and memory CD4^+^ T cells, which have previously undergone activation. Although closely related, there are transcriptional and phenotypic differences in how naive and memory CD4^+^ T cells respond to activation ^13–15^, and there is a high level of heterogeneity within T cells. While naive T cells are generally thought to be a uniform population, memory cells comprise several subpopulations including central memory (T_CM_), effector memory (T_EM_) and effector memory cells re-expressing CD45RA (T_EMRA_). These subsets differ in their proliferation capacity and effector potential, as manifested by the secretion of cytokines that drive inflammation ^16–18^. Additionally, a small subpopulation of regulatory T cells (Tregs) controls T cell activation, thus preventing inflammatory reaction. Transcriptionally, these subpopulations form a continuum of phenotypes within T cells ^19^.

Given the dynamic nature of T cell activation and the heterogeneity of CD4+ T cells, we used single-cell RNA-sequencing (scRNA-seq) to map eQTLs across 119 individuals throughout CD4^+^ T cell activation. Using 655,349 high quality single-cell transcriptomes spanning four time points of T cell activation, we defined 38 stable and transient cell populations. We then reconstructed activation trajectories for naive and memory CD4^+^ T cells, which enabled us to identify eQTL effects manifesting at different points in time and across different subpopulations of cells. We identified 6,407 genes with evidence of genetic regulation, of which 2,265 exhibited dynamic genetic regulation i.e. eQTL effects which changed as a function of time during T cell activation. Finally, we used GWAS summary statistics for eleven immune-mediated diseases and identified 127 genes for which we detected colocalizing signals between eQTLs and GWAS loci. Colocalizing genes were enriched in time-dependent eQTLs, as well as in biological pathways controlling T cell activation. Together, our data suggest that dysregulation of gene expression dynamics during T cell activation could be a key mechanism underlying immune disease and emphasise the importance of accounting for context specific gene expression regulation in interpretation of GWAS signals.

## Results

### Transcriptional response of CD4^+^ T cells to activation at single-cell resolution

To investigate the gene expression changes induced by CD4^+^ T cell activation, we isolated naive and memory CD4^+^ T cells from 119 healthy individuals (**Supplementary Table 1** and **Supplementary Figure 1**) and stimulated the cells using anti-CD3/anti-CD28 coated beads (**Figure 1A** and **Methods**). We profiled gene expression using droplet-based scRNA-seq ^20^ in activated and resting state. We chose three time points to capture cells before (16h) and after the first division (40h), as well as after cells have acquired an effector phenotype (5d) ^19^. This resulted in a high quality transcriptomic data from 655,349 cells (**Methods** and **Supplementary Figure 2**).

**Figure 1.**
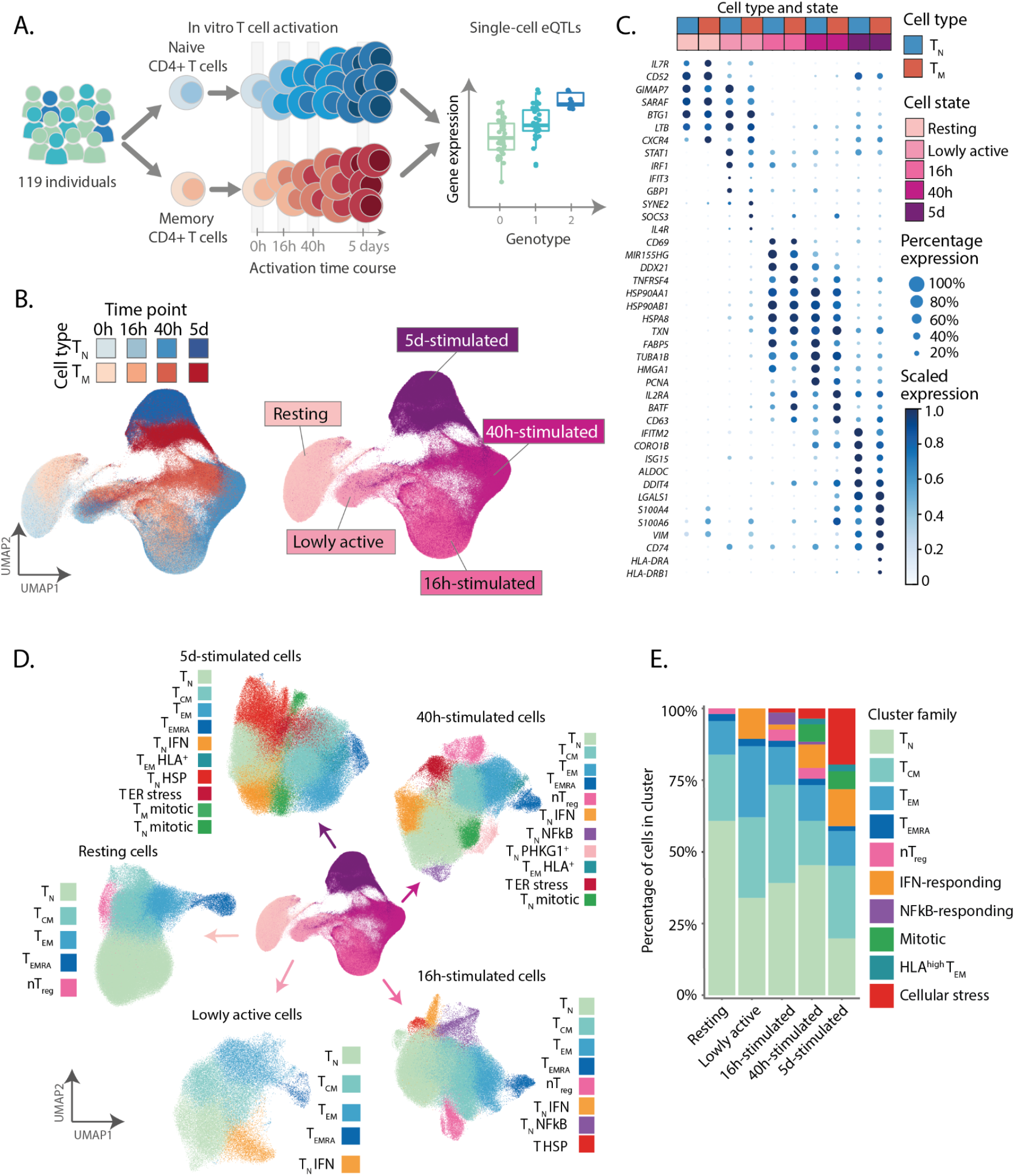
A single-cell transcriptional map of CD4+ T cell activation. **A)** Schematic of the study design. **B)** UMAP embedding of scRNA-seq data for unstimulated CD4^+^ T cells and at three timepoints after activation. Colours represent cell types (blue for T_N_ and red for T_M_) and shades of colours indicate time points (lighter shades for early time points and darker shades for late time points). Right panel represents the five broad cell states. **C)** Dotplot of highly variable gene expression throughout T cell activation. Shades of blue represent average expression in each cell population, and dot sizes represent the proportion of cells expressing the gene. **D)** Separate UMAP embeddings for the five broad cell states. Colours represent cell populations derived from unsupervised clustering. **E)** Proportion of different cluster groups present at each time point. Cell populations defined from clustering were classified into one of 10 families, represented in different colours.

To identify gene expression and cellular composition changes induced by activation, we performed dimensionality reduction and embedding using the uniform manifold approximation (UMAP) ^21^ (**Methods**). Cells separated by time point of stimulation, and formed a gradual progression from resting cells to the most activated cells (largely represented by cells collected at 5d) (**Figure 1B**). This progression was accompanied by changes in known T cell activation markers. For example, an early activation marker, *CD69*, was upregulated at 16h but downregulated at later time points. In contrast, expression of *IL2RA*, a marker of late activation, peaked at 40h and remained present at 5d (**Figure 1C**). We identified a large population of cells that localized between the resting and 16h-stimulated cells (**Figure 1B**). The majority of cells in this group (74%) originated from 16h activated cells, and a proportion (26%) from the 40h time point. We hypothesised that this group represented an early cell state through which cells transitioned as they became activated. To test this, we analysed cells from the 16h and 40h time points independently. We confirmed that at each of these time points cells separated into two clear groups, one of which corresponded to the early transitional state (**Supplementary Figure 4**). Cells in this group expressed 4-fold fewer genes compared to the other cells at their respective activation time points and showed lower expression of a set of CD4+ T cell activation markers which we previously defined ^19^ (**Supplementary Figure 4**). Furthermore, these cells expressed a unique profile characterised by high expression of *STAT1*, *IFIT3*, and *GBP1*, which differed from resting or fully activated cells (**Figure 1C**). Therefore, we concluded that these cells represent a distinct, early activation state which we annotated as lowly active cells.

We next used unsupervised clustering to map the dynamic changes in cell subsets and states throughout the activation time course. This revealed a total of 51 cell clusters, which were merged into 38 cell populations based on their correlated patterns of gene expression (**Supplementary Figure 5** and **Methods**). This included 25 stable subpopulations, which were consistently detected at multiple time points, and 13 transient cell states (**Figure 1D** and **Supplementary Table 2**). Stable subpopulations belonged to one of five phenotypes: naive (T_N_), central memory (T_CM_), effector memory (T_EM_), effector memory re-expressing CD45RA (T_EMRA_), and regulatory (nT_reg_) CD4+ T cells (**Figure 1D and 1E**). The memory T cell pool consists of an average of 59% T_CM_, 30% T_EM_, 5% Tregs, and 5% T_EMRA_. We observed that the percentage of T_EM_ cells decreased with the individual’s age, with a corresponding increase in T_CM_ and T_EMRA_. The distribution of subpopulations was not different between males and females (**Supplementary Figure 7)**. Finally, as we previously demonstrated, these subpopulations formed a naive-to-memory transcriptional progression ^19^.

In contrast to stable subpopulations, transient cell states were only detected at specific activation time points (**Figure 1D and 1E**). For example, we observed a population of cells expressing high levels of interferon (IFN)-induced genes (eg. *IFI6*, *IFIT3*, *ISG15, MX1*) which emerged during early activation (**Supplementary Figure 6**). This is consistent with IFN-induced genes being upregulated during early stages of immune response and their role in the initiation of an inflammatory response. Another transient subpopulation expressed high level of NFκB response genes (eg. *NFKBID*, *REL*, *BCL2A1*) (**Supplementary Figure 6**) and was most dominant at mid-stages of activation, while absent altogether at later stages. Additionally, we observed a population of cells undergoing mitosis, as well as a group of cells expressing high levels of heat shock protein (HSP) family members (e.g. *HSPA1A*, *HSPA1B*, *DNAJB1*; **Supplementary Figure 6**). Both of these were present only during late activation. Interestingly, HSPs have recently been implicated in controlling T cell responses to fever ^22^ We also observed a subset of T_EM_ cells which upregulated MHC class II molecules (eg. *HLA-DRA*, *HLA-DPA1*, *HLA-DRB1*) during late activation (**Supplementary Figure 6**), in agreement with previous descriptions of activation-induced changes in expression dynamics at the HLA locus ^23^. Importantly, individuals uniformly contributed to each cluster, with more variability observed in T_EMRA_ as previously described ^17,24^ (**Supplementary Figure 6F**).

### eQTL map of CD4^+^ T cell activation time course

To study the genetic regulation of gene expression underpinning T cell activation, we performed cis-eQTL mapping. For each time point, we first reconstructed average gene expression profiles per cell type and per individual (i.e. pseudobulk transcriptomes) corresponding to the T_N_ and T_M_ CD4+ T cell types (**Methods**). In total, we detected between 1,545 and 3,006 genes with significant *cis* eQTL effects (eGenes) at different activation time points (**Figure 2A**). We observed that between 210 and 640 eGenes were only detected in individual cell states (**Figure 2B**). For example, kinase *NME4* and purinoceptor *P2RX4* were only detected as eGenes in memory T cells at 16h and 40h of activation, respectively (**Figure 2C**). To assess the extent of this phenomenon and quantify the degree of eQTL sharing across cell types and cell states, we used the multivariate adaptive shrinkage (mashR) method^25^. This analysis showed a higher level of eQTL sharing across cell types within the same time point (**Supplementary Figure 9A**) than across different time points, suggesting that eQTL effect sizes might change throughout T cell activation. Finally, to understand which cell functions might differ in their genetic regulation across cell types, we tested for pathway enrichment in eGenes detected in one cell type and not the other, i.e. in memory but not naive T cells and *vice versa* (**Figure 2D**). To confidently capture a set of genes that are cell-type specific, we only retained genes with an adjusted p-value lower than 0.01 in one cell type, but higher than 0.1 in the other. This showed that, while eGenes specifically detected in 40h-stimulated naive cells were enriched for cell cycle and cell division, eQTLs specific to 16h-stimulated memory cells were enriched in pathways driving mitochondrial organization.

**Figure 2.**
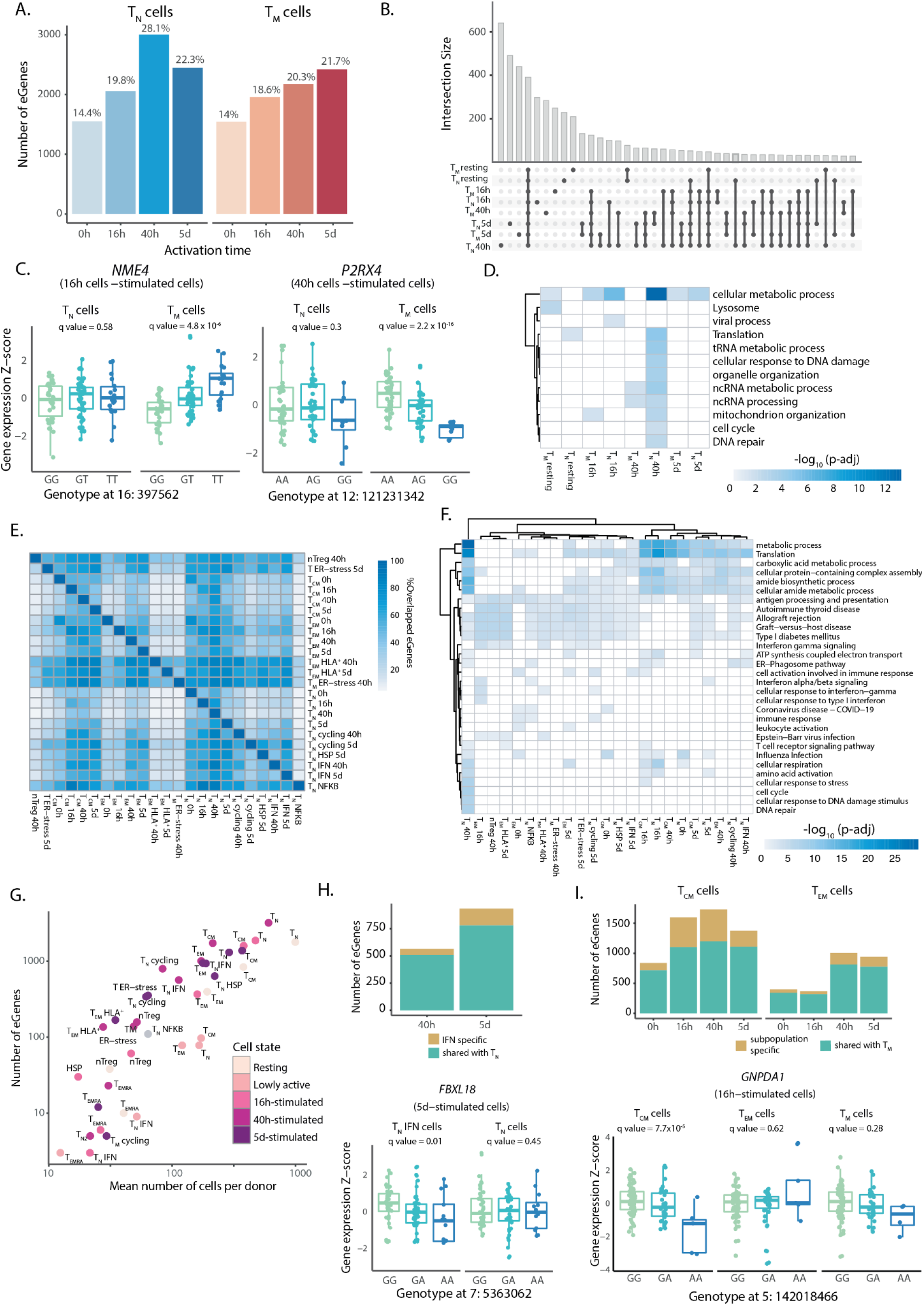
eQTL mapping in resting and activated CD4+ T cells. **A)** Number of significant eGenes detected at each activation time point. Colors represent cell types (blue for T_N_ - naive and red for T_M_ - memory). **B)** Number of significant eGenes shared between cells sampled at each time point. **C)** Example of T memory cell specific eQTLs detected at 16h and 40h. Box plots show the mean expression value of the gene in each sample (Z-scored), stratified by genotype. Each dot represents a measurement obtained from a separate individual. **D)** Enriched pathways in cell state specific eGenes. Color represents adjusted p-value from the enrichment test. **E)** Pairwise comparison of eGenes shared between cell subpopulation. Only subpopulations with more than 100 eGenes were used in this analysis. Green represents the percentage of shared eGenes, yellow represents the percentage of cell type specific eGenes. **F)** Pathways enriched by eGenes detected in each subpopulation. Only subpopulations with more than 100 eGenes were used in this analysis. Colors represent adjusted p-values from the enrichment test. **G**) Scatter plot showing the correlation between number of cells per donor and number of detected eGenes in each cluster. **H)** Number of subpopulation specific eQTLs detected in T_CM_ and T_EM_ cells. Bar plots indicate the numbers of eGenes detected in the T_CM_ and T_EM_ subpopulations that are shared with memory T cells as a whole. Boxplots show an example eQTL specific to the T_CM_ subpopulation. Each dot represents a measurement obtained from a separate individual. **I)** Subpopulation specific eQTLs detected in IFN-responsive clusters. The bar plot indicates the number of eGenes detected in the IFN-responsive subpopulation that are shared with naive T cells as a whole. The boxplots show an example eQTL specific to this subpopulation. Each dot represents a measurement obtained from a separate individual.

To gain a more granular view of gene expression regulation throughout T cell activation we next mapped eQTLs in each of the 38 cell populations that we defined at the single cell level (**Figure 1**). We detected up to 3,197 eGenes per T cell population. As expected, we observed a high overlap between eGenes detected in different T cell clusters (**Figure 2E**). Nevertheless, effector memory cells (T_EM_) and effector memory cells expressing HLA genes (T_EM_ HLA^+^) had a higher number of specific eGenes (62-97%) compared to other subpopulations, suggesting that they are more transcriptionally different than other T cell subsets. eGenes detected in subpopulations were enriched in the immune relevant pathways. For example, in activated Tregs, and various effector T cell subsets we observed enrichment in immune disease genes, IFN-γ signalling and antigen presentation (**Figure 2F**). We also noticed that smaller subpopulations, such as T_EMRA_, yielded a low number of eGenes (3-23) which reflected that the statistical power to detect eGenes was highly correlated with the number of cells profiled per individual per subpopulation (R^2^ = 0.82, p = 4.8×10^10^) (**Figure 2G**). In addition, when we subsampled cells from a cell population and performed eQTL mapping we observed that eGene discovery increased proportionally to the number of analysed cells (**Supplementary Figure 9B**). These results imply that the uncertainty in gene expression estimation from low numbers of cells is high, thus reducing our power to map eQTLs. Despite this limitation, we identified eGenes manifesting in effects in subpopulations and absent in the pseudobulk analysis of the whole naive or memory T cell populations. For example, we identified between 56 and 153 eGenes (10 to 16% of the eGenes detected at the respective time points) which were found in the subpopulation of naive T cells with high levels of IFN-induced genes, but absent in the pseudo-bulk analysis from activated naive T cells (**Figure 2H**) (e.g. *FBXL18)*. Similarly, we identified 47-528 (13-31%) eGenes which were detected in either of the two largest memory cell subpopulations (T_CM_ and T_EM_) but not in pseudo-bulk memory T cells (**Figure 2I**). One example of such a gene is enzyme *GNPDA1,* which is only detected as an eGene in T_CM_ but not in T_EM_ or pseudo-bulk memory cells (**Figure 2I**).

### eQTLs are enriched in proliferation and immune response gene modules

We next sought to understand which transcriptional programmes shape the T cell response throughout the course of activation, and whether eGenes regulate specific cell functions. We computed pairwise gene expression correlations of 11,130 highly expressed and variable genes across 106 individuals and the 38 identified cell populations (**Supplementary Figure 8 and Methods**). This co-expression network captures gene expression patterns at a subpopulation resolution ^26^. Our analysis revealed 12 different gene modules which represent key cellular functions involved in T cell activation (**Figure 3B** and **Supplementary Table 3**). For example, module 4 represented a group of genes involved in the regulation of cell cycle checkpoints and DNA repair mechanisms, and was highly expressed at 40 hours and five days after activation, confirming that the first cell division happens around two days after activation ^27^. Further, module 11 included genes whose expression peaked in lowly active and 16h-stimulated cells and remained high at later activation time points. These genes were involved in IFN-induced antiviral mechanisms such as OAS and ISG15-signalling. These pathways are induced rapidly upon viral infection. Finally, module 2 involved NOTCH4, RUNX2 and RUNX3-signaling. Genes in this module were lowly expressed in resting and lowly active cells and peaked at 16h and 40h, after which genes were downregulated, returning to their baseline levels. This pattern of transient upregulation was also observed for genes involved in RNA metabolism (gene modules 5 and 8).

**Figure 3.**
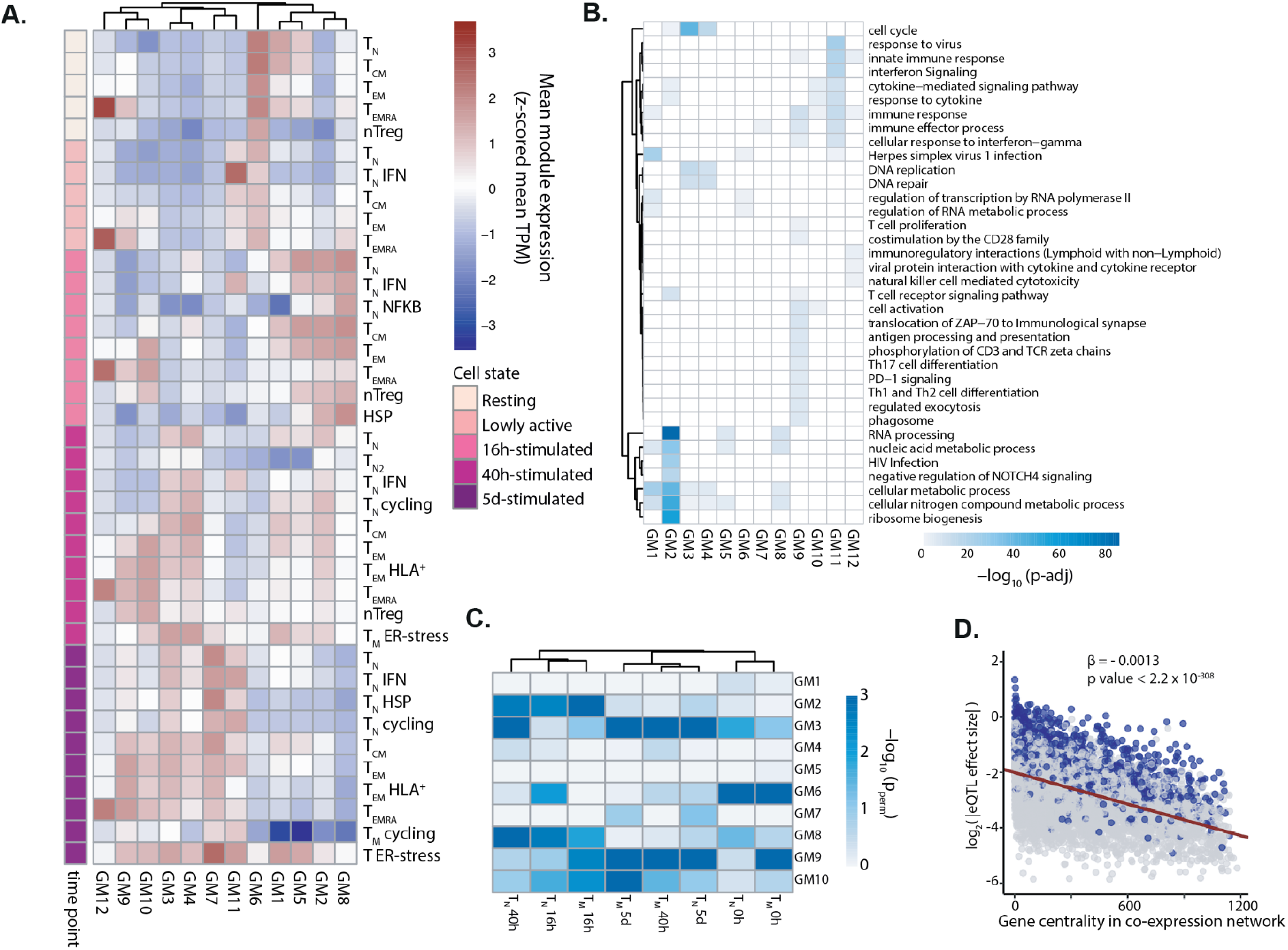
eQTLs are enriched in proliferation and immune response gene modules. A gene co-expression network was built using WGCNA to identify gene modules. **A)** Heatmap showing the expression pattern of the 12 identified gene modules. Rows correspond to cell subpopulations. Colours represent the scaled (Z-scored) average expression of all genes belonging to a module in a given subpopulation. **B)** Pathways enriched in each gene module. Shades of blue represent the log10-transformed enrichment p-values. **C)** Enrichment of eGenes in gene modules. Shades of blue represent the log10-transformed permutation p-value. **D)** Relationship between a gene’s connectivity and the effect size of its lead eQTL variant. All eQTL effect sizes were log2-transformed. Blue dots represent significant eGenes, while gray dots represent genes which do not pass the multiple testing correction.

In addition to separating genes by their temporal dynamics, the co-expression networks also highlighted subpopulation specific gene expression modules, which correspond to effector T cell functions. For example, genes involved in cytokine secretion and interleukin signaling were highly expressed in T_EM_ and T_EMRA_, but not T_CM_ or T_N_ cells (**Figure 3A and 3B**), reflecting the potential of T_EM_ and T_EMRA_ cells to initiate a fast and robust response ^18,19^. This unique property of T_EM_ and T_EMRA_ to mediate a quick effector response upon activation was additionally demonstrated by upregulation of genes within the T cell receptor signaling pathways (i.e. targets of ZAP-70 and downstream of CD3 zeta chain phosphorylation) at an earlier stage of activation than other subpopulations. While T_EM_ and T_EMRA_ showed high expression of the TCR-induced module in the lowly active and 16h states, other subpopulations did not express these genes until 40h after stimulation (**Figure 3A and 3B**). This reinforces the notion that T_EM_ and T_EMRA_ cells respond to stimulation faster than T_CM_ or T_N_ cells. Furthermore, we observed that module 12, which included genes important for cytotoxic function and chemokine signalling, was most highly expressed in resting and activated T_EMRA_ cells (**Figure 3A**). This cytotoxic capacity sets T_EMRA_ cells apart from any other T cell subpopulation.

Next, we asked whether eGenes localised to particular regions of the co-expression networks. Using a permutation strategy (**Methods**), we showed that eGenes detected in activated T cells were particularly enriched in gene modules 2 (metabolism), 3 (cell division) and 9 (immune processes) (**Figure 3C**). This suggests that eGenes are involved in T cell division and function. In contrast, eGenes detected in resting cells showed strongest enrichment in gene module six, which was enriched for RNA metabolism and Herpes infection (**Figure 3C**). Finally, we observed that eQTL effect sizes negatively correlated with the centrality values of the corresponding eGenes in the coexpression network, i.e. eGenes with larger eQTL effects were more likely to be less connected in the network (**Figure 3D**). This suggests that genes at the edges of the co-expression network are more tolerant to variability in gene expression.

### Modelling of time-dependent eQTL effects throughout CD4+ T cell activation

T cells undergo profound transcriptional changes during activation and previous studies have shown that eQTLs can be context specific ^9,28^. Therefore, we sought to assess the role of genetic variation on the regulation of gene expression dynamics throughout T cell activation (dynamic eQTLs). We used trajectory inference ^29^ (**Methods**) to derive a T cell activation trajectory, thus allowing us to model activation time as a continuous variable (**Figure 4A**). We verified that the obtained trajectory agreed with the time points profiled experimentally, with the lowest pseudotime values being assigned to resting cells and the highest to cells activated for five days (**Figure 4A**). Furthermore, the temporal dynamics of well defined T cell activation markers such as *IL7R* (reduced expression upon activation), *CD69* (early activation) and *IL2RA* (early and late activation) (**Figure 4B**) followed their expected patterns, confirming that the inferred trajectory accurately reflects known biology. In total, we identified 5,090 genes for which expression changed as a function of pseudotime (**Supplementary Table 4**). For example, *IRF1* and *TOP2A* were respectively downregulated and upregulated at late stages of activation (**Figure 4B**). Furthermore, we confirmed that dynamically regulated genes were enriched in pathways related to T cell activation, such as DNA replication and regulation of cell cycle, mRNA transcription and processing, protein translation, a metabolic switch to support the electron transport chain, signalling downstream of the TCR, signaling by interleukins, PIP-mediated activation of Akt signaling, and expression of target genes from NF-κB (**Supplementary Table 4**). Finally, we observed that memory T cells were characterised by lower pseudotime values compared to naive cells sampled at the same time points. This is a consequence of memory T cells showing a shorter activation path, likely reflecting a faster activation.

**Figure 4.**
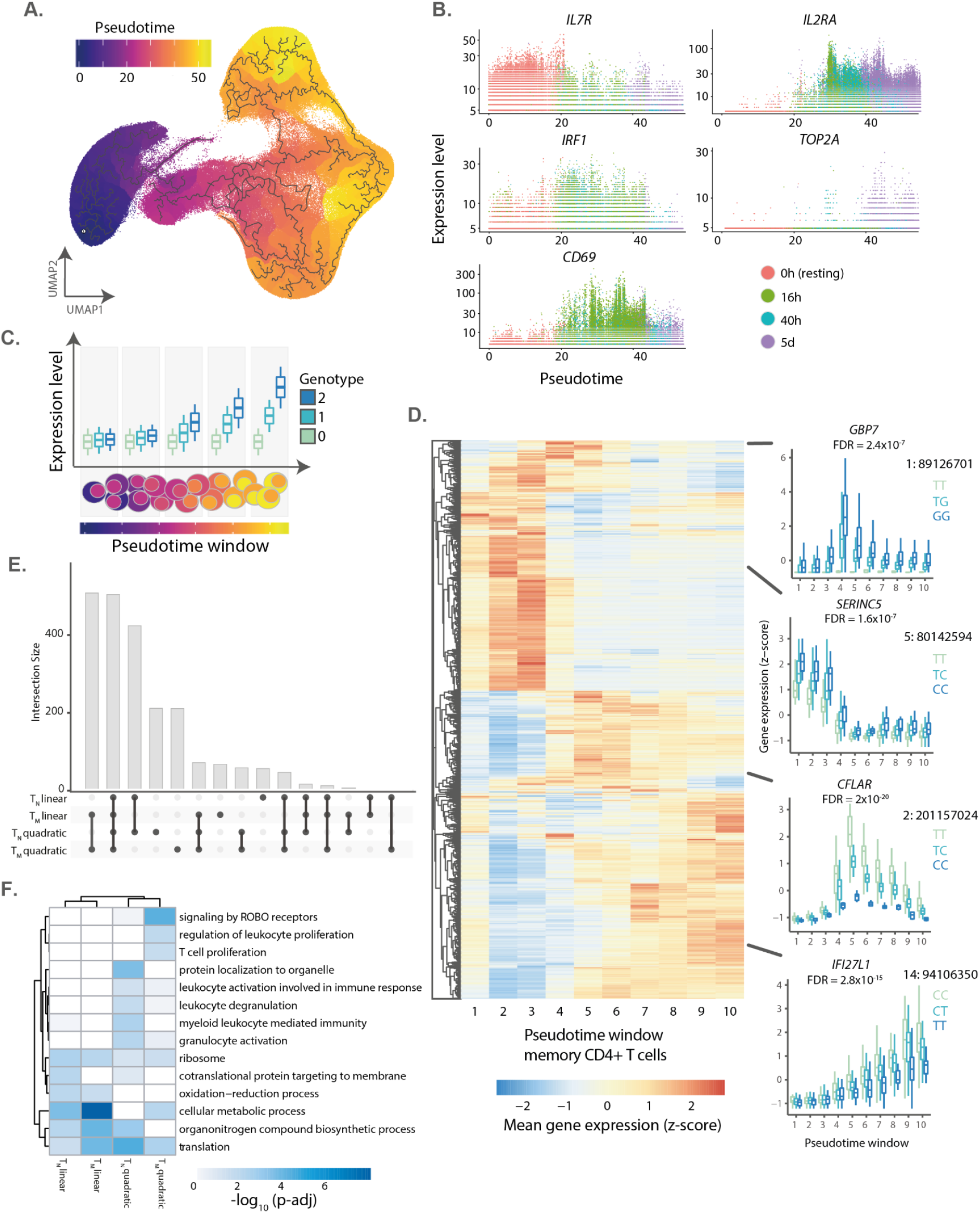
eQTLs with dynamic effects during CD4+ T cell activation. **A)** Cells were ordered into a branched pseudotime trajectory using monocle3. The UMAP embedding shows all cells, coloured by their estimated pseudotime values. Black lines indicate the inferred branched trajectory. **B)** Example genes that significantly change as a function of activation pseudotime. Each dot corresponds to a cell, and colours represent experimental time points. **C)** Schematic of the analysis approach. Cells were split into ten windows of equal cell numbers according to their estimated pseudotime values. Linear and quadratic mixed models were applied to each previously identified eGene to test for an interaction between genotypes and T cell activation pseudotime. **D)** Heatmap showing the expression pattern on each dynamic eGene in memory T cells. Boxplots show examples of non-linear and linear dynamic eQTLs. The average expression of the gene within each pseudotime window was stratified by genotype. **E)** Number of eGenes with evidence of a significant genotype-pseudotime interaction (i.e. dynamic eQTLs) in a linear or quadratic mixed model. **F)** Pathways enriched in linear and quadratic eGenes. Shades of blue represent log10-transformed enrichment p-values.

To model dynamic eQTLs, we divided the inferred pseudotime trajectory into ten bins (i.e. pseudotime deciles) and averaged the expression of genes per individual in each bin (**Methods**). Splitting the trajectory into bins enabled us to control for the numbers of cells and therefore to reliably estimate mean gene expression values. We then applied linear mixed models to test for a significant interaction between genotypes and average pseudotime value per bin (**Figure 4C and Methods**). This enabled us to identify eQTLs for which the effect size changed as a function of activation trajectory. We identified 2,265 genes with dynamic eQTL effects, which comprised 34% of eGenes in our dataset (**Supplementary Table 5**). We applied both linear and quadratic models and observed that most eQTLs followed linear dynamics across the activation trajectory (74% and 76% in naive and memory T cells, respectively) (**Figure 4E**). However, for 502 and 495 genes in naive and memory T cells respectively, we detected a non-linear interaction with activation trajectory. For example, we identified eQTLs with non-linear dynamics for *GBP7* and *CFLAR.* These were only apparent upon activation and the magnitude of their effect sizes peaked at mid stages of the trajectory, significantly diminishing thereafter (**Figure 4D**). In contrast, the magnitude of an eQTL for *SERINC5* peaked at early stages of the trajectory and diminished as cells progressed through activation (**Figure 4D**), as opposed to an eQTL for the interferon alpha inducible gene *IFI27L1* for which an effect size linearly increased along the activation trajectory. Finally, we observed that linear eQTLs were enriched in metabolic pathways, while non-linear eQTLs were enriched both in metabolic processes and immune processes like T cell proliferation and leukocyte degranulation (**Figure 4F**). This suggests that the genetic regulation of immune genes is complex, and evident only during certain stages of T cell activation.

### Colocalization at GWAS loci identifies candidate immune disease genes

The eQTLs mapped in our dataset provide a unique opportunity for functional interpretation of GWAS loci by testing for colocalization between eQTLs and GWAS signals. We used summary statistics for 13 immune-mediated diseases available in the GWAS catalog ^30^ (**Methods**) and tested for colocalization with eQTLs that we mapped in pseudobulk naive and memory CD4^+^ T cells, as well as in the T cell subpopulations and cell states. We used the Bayesian method coloc ^31^ with masking ^32^, which removes the assumption of a single causal variant per locus (**Methods**). We identified 471 unique colocalizations (posterior probability (PP4) > 0.8), corresponding to 247 GWAS loci for 11 diseases and 314 SNP-gene pairs (**Supplementary Table 6**). This enabled us to prioritize 127 candidate disease-causal genes (**Figure 5A**). Importantly, 77 (60%) colocalizing genes were detected only upon activation, and would have been missed by profiling only *ex vivo* cells. Out of those, 47 (37%) were captured specifically in later time points of activation (40h + 5d) (**Figure 5B**). This is important, since eQTL studies mostly relied either on a resting state or a single time point, usually at the early stages of T cell activation ^12,33^.

**Figure 5.**
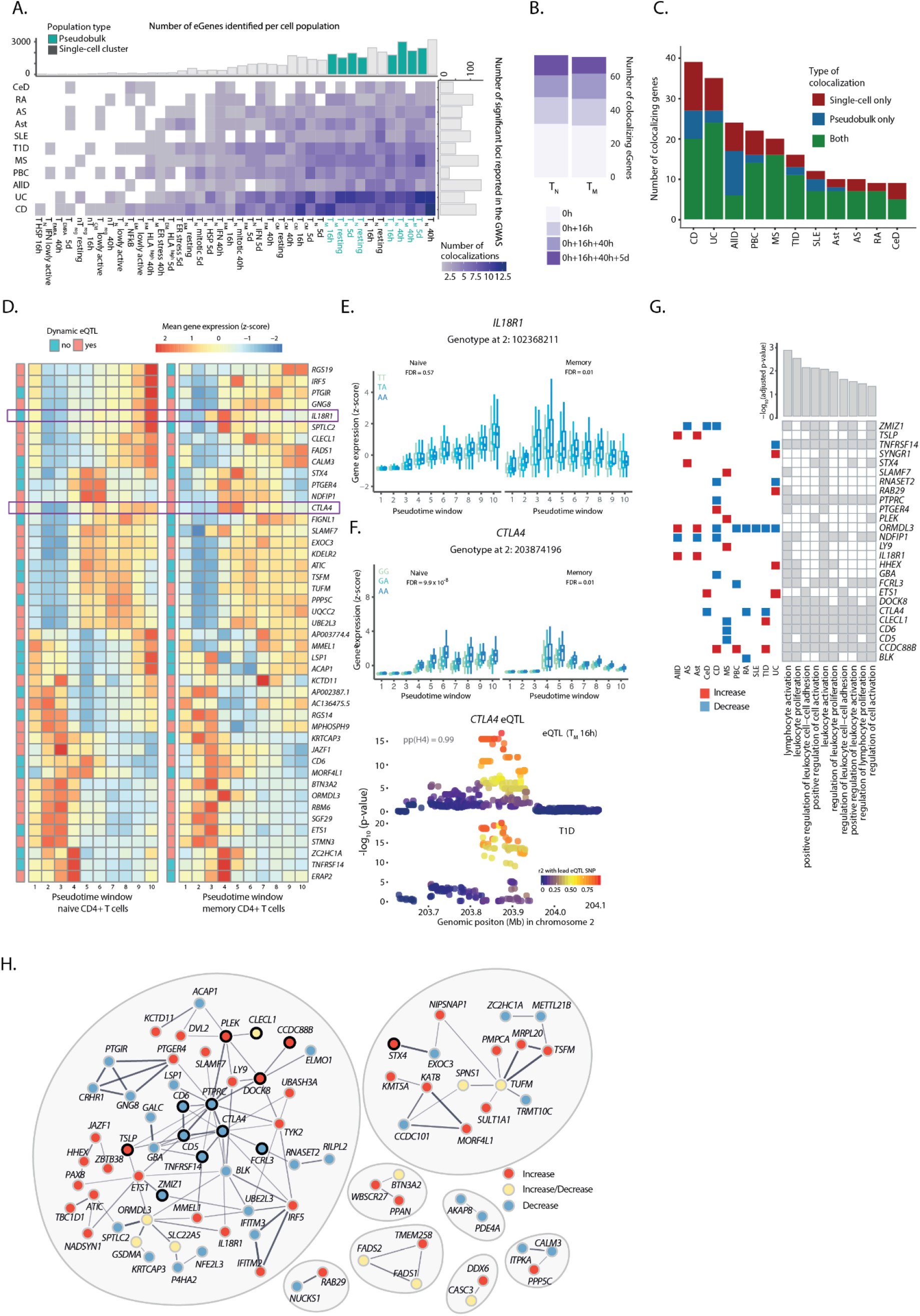
Colocalization of CD4+ T cell eQTLs with GWAS associations for immune diseases. **A)** Number of significant colocalizations between an eQTL and a GWAS signal identified for each cell type-trait combination. Marginal bar plots represent the number of independent associations reported in the GWAS (X axis) and the number of eGenes detected per subpopulation (Y axis). Light and dark bars indicate whole (pseudobulk) cell populations and subpopulations, respectively. **B**) Number of additional colocalizing genes detected in stimulated cells. **C)** Number of colocalizing genes observed in whole cell populations (pseudobulk), subpopulations or both. **D)** Heatmap showing the expression pattern of coloc eGenes in naive and memory T cells. The color of annotation boxis shows genes that are dynamic and static eQTLs. **E)** Boxplot shows *IL18R1* dynamic eQTL. The average expression of the gene within each pseudotime window was stratified by genotype. **F)** Boxplot shows *CTLA4* dynamic eQTL. The average expression of the gene within each pseudotime window was stratified by genotype. Locus plot for a colocalization between a *CTLA4* dynamic eQTL and a GWAS association for type 1 diabetes. Each dot represents a variant, with colors indicating their LD with the lead eQTL variant. **G)** Tile plot shows enriched pathways within colocalizing genes as well as genes driving the enrichment. Barplots show adjusted p-values from the enrichment test. Rectangles on top show the disease that each gene colocalizes with. Red indicates that the disease variant increases the gene expression and blue indicates that it decreases gene expression. **H)** STRING network of colocalizing genes. Red indicates that the disease variant increases the gene expression, blue that it decreases, and yellow that the effect on gene expression is disease dependent. Black outline highlights genes belonging to the top enriched pathway (GO.0050867: positive regulation of cell activation).

Generally, we observed more colocalizations in larger cell populations (for which we were more powered to detect eQTLs) and in traits for which a larger GWAS was available (**Figure 5A**). The traits with the highest number of colocalisations included Crohn’s disease (CD) and ulcerative colitis (UC), followed by allergic diseases (AllD), and previous studies have demonstrated a role of T cells in disease pathobiology ^11,12,34^. However, we also observed that type 1 diabetes (T1D) and systemic lupus erythematosus (SLE), although characterised by a similar number of loci, differed in the proportion of identified colocalizations. In particular, we observed a lower number of colocalizations with SLE variants, in line with the studies pointing towards B cells as the drivers of this disease ^11,35^. We found that 72% of genes colocalized only with one trait, 14% colocalized with two traits and 14% colocalized with 3 or more diseases (**Supplementary Figure 10A**). Overall, we found that 220 disease loci (89%) regulated a single gene, while 22 (9%) and 5 (2%) loci regulated two and three genes in the associated regions, respectively.

While most colocalizing genes were identified in bulk cell types (median per trait = 66%), we observed between 2 and 15 new candidate genes per disease (median per trait = 25%) which could only be detected in individual subpopulations (**Figure 5C**). For example, we identified an eQTL for *TYK2* specifically detected in 16h-stimulated T_EM_ cells colocalized with a CD association (**Supplementary Figure 10B**). Similarly, we identified a colocalization between a CD locus and the *ZMIZ1* eQTL specific to 16h-stimulated T_CM_ cells (**Supplementary Figure 10C**). A closer analysis of *ZMIZ1* revealed that this eQTL is absent in other memory T cell populations such as T_EM_ cells. This causes the signal to be masked in bulk memory cells, where it is no longer detectable (**Supplementary Figure 10C**). Both of these colocalizations are subpopulation and time point-specific, which highlights the importance of measuring gene expression regulation with cell type and cell state resolution to accelerate interpretation of disease associations. We observed no differences in the network connectivity of colocalizing genes compared to the remaining eGenes (**Supplementary Figure 10D**).

Given that the majority of colocalizations were detected in activated T cells we next asked if the expression of these genes showed temporal dynamics. We observed that dynamic eQTLs were significantly enriched in colocalizing eGenes in both naive and memory T cells (36/73 and 44/72 colocalizing genes in naive and memory cells, Fisher’s test p-values 7.9 x 10^−5^ and 2.6 x 10^−7^, respectively). Next, we investigated the gene expression patterns between eGenes that colocalized in both naive and memory cells. While we observed broadly similar gene expression patterns between naive and memory T cells (**Figure 5D**), we also identified genes with differences in expression patterns and genetic regulation in naive and memory T cells. For example, IL-18 receptor (*IL18R1*) is a dynamic eQTL only in memory T cells, and it is highly expressed during early activation of memory T cells, while the expression peaks at late activation in naive T cells (**Figure 5E**). In another example, we observed that *CTLA4*, which was detected as a dynamic eQTL in both memory and naive T cells, showed different regulation in the two cell types (**Figure 5F**); in naive cells upon activation cells upregulated and maintained high expression of *CTLA4*, while in memory cells the expression peaked at early activation and then diminished. This eQTL colocalized with a type 1 diabetes associated locus and individuals carrying the disease risk allele showed lower expression of *CTLA4*. This example is particularly illustrative because of the role *CTLA4* plays in regulating the immune response. Reducing expression of *CTLA4* at early stages of cell activation could result in reduced ability to suppress T cell activation (as *CTLA4* is a key mediator of this process) and would contribute towards excessive T cell activation in disease settings. Additionally, the same *CTLA4* eQTL variant colocalized with suggestive association signals for rheumatoid arthritis and celiac disease, in fact, *CTLA4* antibody (e.g. abatacept) is used as a treatment in rheumatoid arthritis (**Supplementary Table 6**).

Finally, we assessed whether immune disease loci affected any specific cellular functions. We observed that, collectively, colocalizing genes were significantly enriched in pathways involved in the regulation of T cell activation and proliferation (**Figure 5G**). There were 26 genes driving this enrichment, and mainly included the colocalizing genes with association signals shared across two or more diseases. For 24 out of 26 genes the direction of effect of the risk allele on gene expression was consistent between traits. Most of the colocalizing genes clustered into connected modules based on the STRING database ^36^, i.e. the genes were physically interacting at the protein level or were co-expressed across a broader range of tissues (**Figure 5H**). Furthermore, neighbouring genes within these modules tended to be perturbed in the same direction by immune disease variants. For example, we observed a large module of interconnected genes, 12 of which were involved in the regulation of T cell activation and proliferation. For example, *PTPRC* was directly connected to *CD6*, *CD5*, *CTLA4*, and *TNFRSF14*. Strikingly, all of these genes were downregulated by risk alleles at nearby disease loci, suggesting that reduced expression of these genes may drive the disease development. Together, our results demonstrate that immune disease loci colocalize with genes that are involved in the regulation of T cell activation, and that genes with similar functions tend to be modulated by disease risk alleles in the same direction.

## Discussion

T cell activation plays a critical role in regulating the immune response, and dysregulation of this process results in poor response to infections, development of common inflammatory and autoimmune diseases, and in severe instances primary immunodeficiencies. Our study provides a high-resolution dynamic map of the transcriptional changes which accompany cells’ transition from a resting state through to three time points of activation.

By using single-cell transcriptional profiling across 655,349 CD4^+^ T cells we provide an unbiased view of the T cell response to stimulation, detecting a variety of cell states which change throughout activation. We identified 38 different cell states, including 13 transient cell states. The granularity of the single-cell resolution in our study provides new explanations to previous results from bulk gene expression. For example, a previous study of CD4^+^ T cell activation concluded that T cells upregulate a module of IFN-related genes early upon TCR engagement ^37^. Here we recapitulate this observation and further resolve it to one specific subpopulation of naive cells, rather than the whole T cell pool. We also demonstrate that previously described changes in the expression of class II MHC molecules upon T cell activation ^23^, are more precisely driven by T_EM_ cells. Therefore, our data provides a unique resource for the interpretation of existing and design of future studies of T cell function.

Often eQTLs obtained from bulk RNA-seq mask cell-type specific effects ^38^, but such gene expression effects can be mapped with single-cell transcriptomics ^39^. The available immune cell eQTL resources ^33,40^, including those capturing T cell activation ^37^, have relied on sorting cells based on the expression of cell surface markers. While sorting enriches cell populations of certain characteristics, it cannot capture cellular heterogeneity in the sorted populations.

The single-cell approach of our study allowed us to map eQTLs individually within clusters identified in an unbiased manner, providing insights into cellular processes that are genetically regulated across different cell types. For example, we observed that at 40h of activation eGenes detected in naive CD4^+^ T cells (T_N_ 40h) were particularly strongly enriched in different cell metabolic processes, T cell receptor signalling and cell cycle pathways. These profiles set the T_N_ 40h cell state firmly apart from other cell states. On the other hand, we observed that eGenes detected in more differentiated cell clusters, such as T_N NFκB_, T_regs_, and T_EM_ cells were enriched in pathways attributed to cell effector functions (antigen processing, interferon gamma signalling) and pathways implicated in dysregulation of immune homeostasis in tissues (type I diabetes, autoimmune thyroid disease and allograft rejection). The results of our study will help to infer in detail the effects of genetic regulation on the development of effector T cell functions, and could in the future inform cell engineering approaches.

Expression QTLs can also be context specific, including those resulting from a response to a specific stimulus ^9,28^. However, most existing eQTL resources have profiled cells and tissues at steady states. While these resources have been instrumental in interpreting GWAS signals, the proportion of disease associated variants colocalizing with eQTLs remains low ^41^. By profiling multiple time points of T cell activation, our study further emphasises the complexity of context specific gene expression regulation. Importantly, had we only focused on the resting state, we would have missed the majority of the eQTL effects, impacting our ability to identify eGenes colocalizing with disease associated variants (only 40% in resting state). Indeed, eQTLs colocalizing with disease associated variants were enriched for eGenes with dynamic gene regulation as compared to all the eGenes in our dataset. This emphasizes the challenges of translating GWAS signals to function and could explain why, at present, eQTL colocalizations are explaining only a small proportion of GWAS associations.

Finally, we show that our results capture relevant disease biology and can be used to prioritize novel drug targets. For example, our study found a *CTLA4* eQTL which colocalizes with GWAS associations for three immune diseases, and in all the cases the risk allele decreased gene expression. This agrees with the known role of CTLA4 in removing costimulatory molecules from the surface of antigen presenting cells, thus downregulating T cell activation ^42^. Thus, a partial reduction of CTLA4 function results in impaired immune regulation and higher odds of initiating an autoimmune response ^43^. Furthermore, this is in line with an existing therapeutic approach in which a CTLA4 fusion protein is administered to patients with rheumatoid arthritis to regulate T cell activation and help reduce inflammation ^44^. Importantly, we show that the expression of *CTLA4* is dynamically regulated, peaking during early cell activation.

Similarly, a *TYK2* eQTL, for which we detected the effect in T_EM_ cells at 16 hours of activation, colocalizes with variants associated with Crohn’s disease. The *TYK2* locus is associated with 10 different autoimmune disorders with at least three independent signals reported ^1,45,46^. One of these signals is explained by a missense variant, which reduces signalling downstream of several cytokine receptors, resulting in a protective effect for immune diseases ^1^. Here, we show a similar effect, where individuals carrying a protective allele for Crohn’s disease have lower expression of *TYK2* in T_EM_ cells at 16 hours of activation. Inhibition of TYK2 in treatment of inflammatory diseases is currently in clinical trials ^47^. In addition, a *SLAMF7* eQTL detected at 40h of activation colocalizes with a GWAS signal for MS, and SLAMF7 regulators are used in the treatment for multiple myeloma ^48^. Therefore, MS could be an alternate indication. These examples imply that colocalizing genes from our study could have therapeutic value for immune diseases.

One limitation of this study is that we investigated T cell activation in healthy individuals. As the disease does not alter the underlying genetics, we can identify eQTLs that determine immune disease susceptibility. However, we are likely missing T cell states specific for inflammatory conditions or eQTL signals that only occur following the disease onset. Future studies in disease cohorts will be required to understand the differences in eQTL discovery before and after onset of inflammatory diseases.

Our results emphasize the importance of mapping context-specific gene expression regulation, provide insights into the mechanisms of genetic susceptibility of immune diseases, and help prioritize new therapeutic targets.

## Acknowledgements

We also thank all the donors that participated in this study. We thank the Wellcome Sanger Institute Flow Cytometry facility for their assistance in cell sorting, the Sequencing facility and Cellular Genetics Informatics team for their contribution to data generation and processing. We thank Ian Dunham and Leland Taylor for critical feedback on the manuscript. This work was funded by the Open Targets grant (OTAR040) awarded to G.T. G.T. is supported by the Wellcome Trust (grant WT206194). E.C.-G. is supported by a Gates Cambridge Scholarship (OPP1144).

## Methods

### Cell isolation and stimulation

Blood samples were obtained from 119 healthy individuals of British ancestry. Of these, 67 were male (53.7%) and 52 female (56.3%), and the mean age of the cohort was 47 years (sd = 15.61 years) (**Supplementary Figure 1A**). Human biological samples were sourced ethically and their research use was in accord with the terms of informed consent under an IRB/EC approved protocol (15/NW/0282).

Peripheral blood mononuclear cells (PBMCs) were isolated using Ficoll-Paque PLUS (GE healthcare, Buckingham, UK) density gradient centrifugation. Naïve (CD25- CD45RA+ CD45RO-) and memory (CD25- CD45RA- CD45RO+) CD4+ T cells were isolated from the PBMC fraction using EasySep® naïve CD4+ T cell isolation kits and memory CD4+ T cell enrichment kits (StemCell Technologies, Meylan, France) according to the manufacturer’s instructions. Naive and memory T cells were then stimulated with anti-CD3/anti-CD28 human T-Activator Dynabeads® (Invitrogen) at a 1:2 beads-to-cells ratio. Cells were harvested after 16 hours, 40 hours, and 5 days of stimulation. In addition, unstimulated cells kept in culture without any beads for 16 hours were used as a negative control (i.e. zero hours of activation).

### Single-cell RNA-sequencing

Upon harvesting, cells were resuspended in RPMI media to obtain a single-cell suspension. Next, cells were stained with the live/dead dye 4’,6-diamidino-2-phenylindole (DAPI) and dead cells were removed from the suspension using fluorescence-activated cell sorting (FACS). Live cells were resuspended in phosphate buffer saline (PBS), at which point cells obtained from different individuals but belonging to the same experimental condition were mixed together at equal ratios to form a single cell suspension (i.e. pool). Each pool corresponded to a mix of cells from four to six different individuals (median = 6), and we processed a total of 172 pools.

Cells were next processed for single-cell RNA-sequencing using the 10X-Genomics 3’ v2 kit ^20^, as specified by the manufacturer’s instructions. Namely, 1 x 10^4^ cells were loaded into each inlet of a 10X-Genomics Chromium controller in order to create GEM emulsions. Each experimental condition was loaded in a separate inlet. The targeted recovery was 6,000 cells per pool. Reverse transcription was performed on the emulsion, after which cDNA was purified, amplified, and used to construct RNA-sequencing libraries. These libraries were sequenced using the Illumina HiSeq 4000 platform, with 75 bp paired-end reads and one cell pool per sequencing lane.

### Genotyping

Genomic DNA was isolated from a suspension of 1 x 10^6^ PBMCs from each individual in the study using a DNA isolation kit (Qiagen). Genotyping was then performed using the Infinium CoreExome-24 (v1.3) chip (Illumina).

### Genotyping data analysis and imputation

Quality control was performed by removing samples with <95% called genotypes, as well as keeping only variants with MAF > 5%, SNP call rate > 95%, and in Hardy-Weinberg equilibrium (HWE; p-value < 0.001).

Imputation of untyped variants was performed as described in Bossini-Castillo et al ^49^. Briefly we used BEAGLE 4.1 ^50^ with a reference panel consisting of the 1000 Genomes Phase 3 ^51^ and the UK10K samples ^52^. Variants derived from imputation were quality filtered using the following parameters: allelic R-squared (AR2) >= 0.8, HWE p-value < 0.001, and MAF > 10%. In total we retained 4,641,747 variants after imputation, which were used for eQTL analysis. We confirmed that all but one of the individuals in our cohort clustered with the British in England and Scotland (GBR) population in PCA space (**Supplementary Figure 1B**). This individual clustered with the Finnish population and was removed from any further analyses. Finally, related individuals were kept for every analysis except eQTL mapping, where one random individual from each pair was used.

### Single-cell RNA-sequencing data analysis

#### Data processing and quality controls

Raw scRNA-seq data were processed using the Cell Ranger Single-Cell Software Suite ^20^ (v3.0.0, 10X-Genomics). In brief, reads were first assigned to cells and then aligned to the human genome using STAR ^53^, with the hg38 build of the human genome (GRCh38) as a reference for alignment. Ensembl (v93) was used as a reference for gene annotation, and gene expression was quantified using reads assigned to cells and confidently mapped to the genome.

Results from RNA quantification in Cell Ranger were imported into Python (v3.8.1) and analysed using scanpy (v1.4.4) ^54^. Samples with less than 70% of reads mapping to cells were discarded. This resulted in 142 (82%) cell pools and 106 (89%) individuals being kept after quality filters. In addition, any cells with less than 200 detected genes, an unusually high number of genes (defined as over four standard deviations above the mean number of detected genes), or more than 10% of reads mapping to mitochondrial genes were removed from the data set. Finally, any genes detected in less than 10 cells were discarded. This resulted in 713,403 cells (96.77% of total) and 23,360 genes passing quality filters.

#### Deconvolution of single cells by genotype

Each scRNA-seq sample comprised a mix of cells from unrelated individuals. Thus, natural genetic variation was used to assign cells to their respective individuals. First, a list of common exonic variants was compiled from the 1000 genomes project phase 3 exome-sequencing data ^51^. This list included any variants with a minor allele frequency of at least 5% in the European population. Next, cellSNP (v0.99) ^55^ was used to generate pileups at the genomic location of these variants. These pileups, in combination with the variants called from genotyping in each individual, was used as an input for Vireo (v1), ^55^. Vireo uses a Bayesian approach to infer which cells belong to the same individual based on the genetic variants detected within scRNA-seq reads. Any cells labelled as “unassigned” (less than 0.9 posterior probability of belonging to any individual) or “doublets” (containing mixed genotypes) by Vireo were discarded. On average, 92% of the cells in each pool were unambiguously assigned to a single individual in the cohort (**Supplementary Figure 2**).

#### Cell cycle scoring

After quality control, the number of unique molecular identifiers (UMIs) mapping to each gene in each single cell were normalized for library size and log-transformed using scanpy’s default normalization parameters ^54^. Next, a publicly available list of cell cycle genes ^56^ was used in combination with scanpy to perform cell cycle scoring and assign cells to their respective stage of the cell cycle.

#### Exploratory data analysis and removal of cellular contaminations

We performed exploratory analysis at each experimental time point independently. Cells collected at the same time point were first loaded into scanpy, where normalised log-transformed UMI counts were used to identify highly variable genes. Between 701 and 1,668 highly variable genes were detected at each time point (mean = 1,301). Only highly variable genes were used as a basis for the remaining analyses in this section.

Having identified highly variable genes, technical covariates (cell culture batch) and unwanted sources of biological variation (i.e. number of UMIs per cell, proportion of reads mapping to mitochondrial genes, cell cycle scores, and reported sex) were regressed out using scanpy’s regress_out() function. Next, log-UMI counts were scaled (setting 10 as the maximum value) and used as an input for principal component analysis. The first 40 principal components were used to build a k-nearest neighbours (kNN) graph (with k=15), which was used as an input for embedding and visualization with the uniform manifold approximation and projection (UMAP) algorithm ^21^. This kNN graph was further used for unsupervised clustering using the Leiden algorithm ^57^.

At this stage, cell clustering revealed a low proportion of three contaminating cell types which were consistently detected at each time point: B cells, CD8+ T cells, and antigen-presenting cells (APCs). Furthermore, two additional sources of contamination (SOX4+ precursor cells, and cells expressing hallmarks of cell culture stress) were detected at zero hours of activation (**Supplementary Figure 3**). Cell contaminations were removed from the data set, resulting in 655,349 (91.86% of total) high quality cells kept and successfully annotated as CD4+ T cells.

#### Identification of a lowly active T cell subpopulation

Having removed cellular contaminations, highly variable genes were recalculated and the analysis described in the previous section (i.e. batch regression, scaling, PCA, graph construction, embedding, and clustering) was repeated using CD4+ T cells only. Cells sampled at 16h and 40h showed a clear separation into two groups, one of which expressed a significantly lower number of genes and showed comparatively lower levels of previously described T cell activation markers ^19^ (**Supplementary Figure 4A**). This population of lowly active cells was separated from its original time point and treated as an independent group for clustering.

#### Clustering and cluster annotation

Unsupervised clustering was applied independently to the five cell groups of cells identified in the study (resting, lowly active, 16 hours, 40 hours, and five days) based on their respective kNN graphs and using the Leiden algorithm ^57^. This resulted in 51 cell clusters. The similarity of these clusters to each other was assessed by performing PCA on the full data set (i.e. all cells) and estimating the Euclidean distance between pairs of clusters (from cluster centre to cluster centre) based on the first 100 principal components. Clusters with high levels of similarity or overlapping biological characteristics were merged together (**Supplementary Figure 5B**). This resulted in 38 distinct groups of cells. Gene markers for each of these groups were identified using scanpy’s built-in function for gene ranking, which uses a T test to compare the average expression of a gene in a cluster versus its expression outside the cluster. Each cell group was annotated by comparing its inferred marker genes with known cell type markers reported in the literature.

#### Ordering of cells in a pseudotime trajectory

To perform trajectory inference, raw gene expression measurements for all CD4+ T cells in the study (i.e. 655,349 cells spanning all time points) were imported into R (v3.6.1) and analysed using monocle3 (v0.2.0) ^29^. As opposed to other analyses, where cells from each time point were treated independently, here some unwanted sources of variation such as cell cycle scores correlated with the biological process of interest (i.e. T cell activation). Thus, we implemented a hierarchical batch regression approach, where cell cycle scores were first regressed within each time point, followed by batch regression in the full data set. In brief, principal component analysis was performed based on all cells using monocle3’s PCA implementation. Next, a matrix containing the first 100 principal component coordinates for each cell was split by time point. Cell cycle effects were then regressed from each sub-matrix independently using limma’s lmFit function ^58^. Finally, these cell cycle-corrected matrices were merged back into a full PCA matrix and cell culture batch effects were regressed based on the full data set using the mutual nearest neighbours (MNN) algorithm ^59^.

After batch correction, the first 100 principal components were used to build a kNN graph and this graph was embedded into a two-dimensional space using UMAP. Finally, UMAP coordinates were used to infer a branched pseudotime trajectory using monocle3’s learn_graph function. To identify genes that changed as a function of pseudotime, monocle3’s graph test was applied to all genes. This test assesses whether cells adjacent in the trajectory show more correlated expression of a gene than cells which are far apart (i.e. autocorrelation). Correction for multiple testing was performed using the Q value procedure ^60^. A gene was considered as significantly associated with pseudotime if it had a Q value ≤ 0.05 and a Moran’s I (a measurement of the magnitude of autocorrelation) larger than 0.05 ^61^.

#### Co-expression network analysis

In order to preserve most of the resolution provided by scRNA-seq while also minimising zero-inflation and technical noise, gene expression was aggregated per individual and per cluster into a pseudobulk count matrix. More specifically, raw UMI counts per gene were summed across all cells belonging to the same cluster and to the same individual. Summed expression values were normalised by library size and gene length, resulting in a single expression matrix with transcript per million (TPM) units (i.e. a pseudobulk matrix).

The resulting pseudobulk matrix was imported into R (v3.6.1) and analysed using the weighted gene co-expression network analysis (WGCNA) package (v1.69) ^26^. Only genes with ≥ 1 TPM in at least 30 samples were used for this analysis. TPM values were first log-transformed, after which unwanted sources of variation (i.e. cell culture batch, reported sex, and age) were regressed out using limma’s (v3.40.6) removeBatchEffect() function ^58^. Next, genes were filtered by their level of variability, with only genes showing a standard deviation ≥ 0.1 across samples being kept. This resulted in 11,130 genes taken forward for network construction.

The functions available in WGCNA were used to calculate network properties for these genes, such as their level of connectivity, as well as to build an adjacency matrix. The soft power parameter was set to 4 in these analyses, based on an evaluation of the [0,20] parameter space. The resulting adjacency matrix was transformed into Topological Overlap Matrices (TOM) of similarity and dissimilarity. Finally, hierarchical clustering was applied to the TOM dissimilarity matrix in order to build a dendrogram of genes. Gene modules were inferred from this dendrogram using R’s dynamicTreeCut package (v1.63.1). Mean module expression values were calculated as the average expression of all genes belonging to that module in a given sample.

### Pathway enrichment analysis

All pathway enrichment analyses were performed using gprofiler2 (v 0.1.9), setting the gene list of interest as an unordered query and using all genes detected in the study as the background. Only enriched terms derived from the Kyoto Encyclopedia of Genes and Genomes (KEGG) or the REACTOME database were kept. gprofiler2’s built-in approach (gSCS) was used for multiple testing correction. Enriched pathways were visualized in R using the pheatmap package (v1.0.12).

### Expression quantitative trait loci (eQTL) mapping

#### Mapping of eQTLs

For each gene we calculated mean expression per cluster per donor. To ensure the high quality eQTL mapping, we only kept genes with non-zero expression in at least 10% of donors and mean count per million (cmp) higher than one. We retained between 8,940 and 11,516 genes. To identify cis-eQTLs we used tensorQTL (v1.0.3) ^62^ to run a linear regression for each SNP-gene pair, using a 500 kilobase window within the transcription start site (TSS) of each gene (i.e. cis_nominal mode). We regressed the first 15 gene expression principal components from this analysis, so as to capture the confounders within our dataset. To correct for the number of association tests performed per gene, we used a cis permutation pass per gene with 1000 permutations. Finally, to correct for the number of genes tested and identify significant eGenes we performed a q-value correction ^63^ for the top associated SNP-gene pair, setting a q-value threshold of 0.1.

#### Analysis of eQTL sharing across cell types

To assess the sharing between eQTLs, we performed a meta-analysis across cell types and cell states using the multivariate adaptive shrinkage (mashR) method ^25^. MashR is a Bayesian method which estimates the pairwise level of sharing between cell states, where an eQTL is defined as shared if it has the same effect size (within a factor of 0.5) and direction in two cell states.

#### Modelling of eQTL effect sizes as a function of network centrality

To assess the relationship between a gene’s genetic regulation and network centrality, the effect size of each gene’s lead eQTL variant was modelled as a function of the gene’s centrality value in the co-expression network described above. This was first done assuming a linear relationship. However, substantial heteroskedasticity was observed, which suggested a non-linear relationship, as confirmed using a Breusch-Pagan heteroskedasticity test ^64^. Thus, we log-transformed the eQTL effect sizes, which resulted in homoskedastic data and a strong linear relationship between the variables. All linear models were built and tested using base R’s lm() function.

#### Modelling of dynamic psudotime-dependent eQTL effects

To identify pseudotime-dependent eQTL effects, we divided the activation trajectory into 10 windows containing roughly equal numbers of cells (i.e. pseudotime deciles) and averaged the expression of each gene per individual within each window. To facilitate the interpretation of coefficients, pseudotime windows were scaled from 0 to 1 prior to this analysis. In order to account for the higher correlation in expression values derived from the same individual at multiple pseudotime windows, we applied linear (1) and quadratic (2) mixed models, with individuals modelled as random intercepts. We used these models to test for a significant interaction between genotypes (i.e. the genetic dosage carried by each individual at the lead eQTL variant for that gene) and pseudotime, as follows:

1. Z_score ∼ Genotype + Pseudotime + Cell_culture_batch + Sex + Age + Genotype*Pseudotime + (1 | Donor)
2. Z_score ∼ Genotype + Pseudotime + Pseudotime^2^ + Cell_culture_batch + Sex + Age + Genotype*Pseudotime + Genotype*Pseudotime^2^ + (1 | Donor)

In both cases, the null model was computed using the same parameters while excluding the Genotype*Pseudotime and Genotype*Pseudotime^2^ terms. P-values were calculated by comparing each model to its respective null model using analysis of variance (ANOVA). All models were implemented in R using the *lmer()* function. In order to reduce the burden imposed by multiple testing, we only applied this approach to variants previously identified as significant lead eQTL variants for a gene by tensorQTL in at least one time point. This was done separately for naive and memory T cells.

### Estimation of pairwise LD

We performed LD calculations based on the individual-level genotype information for the individuals in this study obtained from genotyping. Namely, we used PLINK (v1.90b4) ^65^ to calculate the correlation between pairs of genetic variants across all individuals in the cohort, using either the --r or the --r2 flags. We restricted this calculation to variants with MAF > 0.01, computing correlations for any pairs of variants located within 1 Mb of each other and at most 500 variants away. The command used was: *plink --r/--r2 --ld-window 500 --ld-window-kb 1000 --maf 0.01 --vcf-half-call h*

### Integration of eQTLs with GWAS signals

#### Pre-processing of GWAS summary statistics

Full summary statistics files from previous GWAS studies were downloaded from the GWAS catalogue ^66^. These files corresponded to a recent release containing harmonised summary statistics, which were lifted over to build GRCh38 of the genome and for which the effect size and effect direction of each variant was aligned to the reported alternative allele ^30^. Any signals coming from the X or Y chromosomes, as well as from the MHC region (ch6:28,510,120 – chr6:33,480,577) were discarded. These summary statistics corresponded to 13 immune-mediated diseases: celiac disease ^67^, rheumatoid arthritis ^68^, ankylosing spondylitis ^69^, asthma ^70^, systemic lupus erythematosus ^71^, type 1 diabetes ^72^, multiple sclerosis ^73^, primary biliary cirrhosis ^74^, allergic disease ^75^, juvenile idiopathic arthritis ^76^, psoriasis ^77^, ulcerative colitis, and Crohn’s disease ^78^.

#### Colocalization analysis

Genomic loci of interest were identified by intersecting eQTL signals in each cell type with GWAS loci for 13 immune-mediated diseases. For each trait-cell type pair, we applied colocalization to any locus where a lead variant for a significant eQTL (q value < 0.1) was located within 100 kb and in high LD (R^2^ > 0.5) with a significant GWAS variant (i.e. any GWAS variant with nominal p value < 1 x 10^−5^, which enabled us to capture suggestive association signals). In addition, we required at least 50 variants to be available for testing at each candidate locus.

At each of these loci, coloc (v4.0.4) was used to test for colocalization between the eQTL and the GWAS signals. Importantly, these analyses were based on the recently developed masking approach, which relaxes coloc’s previous assumption of a single causal variant per locus ^32^. This is similar to performing conditional analyses at each locus. In brief, we defined a 500 kb window centred at the lead eQTL variant and tested for colocalization using all common variants located in the window and present in both the eQTL and the GWAS summary statistics. We used the pairwise LD calculations from our cohort as a basis for masking, setting an R^2^ threshold of 0.01 to separate independent signals. Coloc’s prior parameters were set to their recommended values in the most recent publication ^32^ (p1=1×10^−4^, p2=1×10^−4^, p12=5×10^−6^). Significant colocalizations were defined as any instances where the estimated posterior probability of a shared causal variant (PP4) was ≥ 0.8. In order to discard potential false positives due to noisy association signals, we only kept for further analysis traits with more than one significant colocalization (11 out of 13 traits).

To infer the relationship between gene expression and disease risk at each locus, we estimated the GWAS and eQTL effect sizes (i.e. log_e_OR and gene expression Z-score) for the GWAS variant in highest LD with the lead eQTL variant at the locus. We concluded that a variant increased disease risk via an increase in gene expression if the variant had the same direction of effects in both studies. In the opposite case, we concluded that the variant increased disease risk via a decrease in gene expression. If the same variant had different estimates of eQTL effect size in different T cell populations, we required that all effect sizes had the same direction.

### Code and data availability

The raw single-cell RNA-sequencing data described in this study will be deposited in the European Genome-Phenome Archive (EGA) upon manuscript publication. In addition, eQTL lead signals and summary statistics will be deposited in the eQTL catalogue (https://www.ebi.ac.uk/eqtl/). A count matrix with processed scRNA-seq UMI counts for all cells in the study, as well as lower dimensional representations of the data set, will be made available via the Open Targets website. All the codes used for processing and analyzing the data in this study are compiled in a GitHub repository, which will be made publicly available upon manuscript publication.

## Supplementary Figures

**Supplementary Figure 1.**
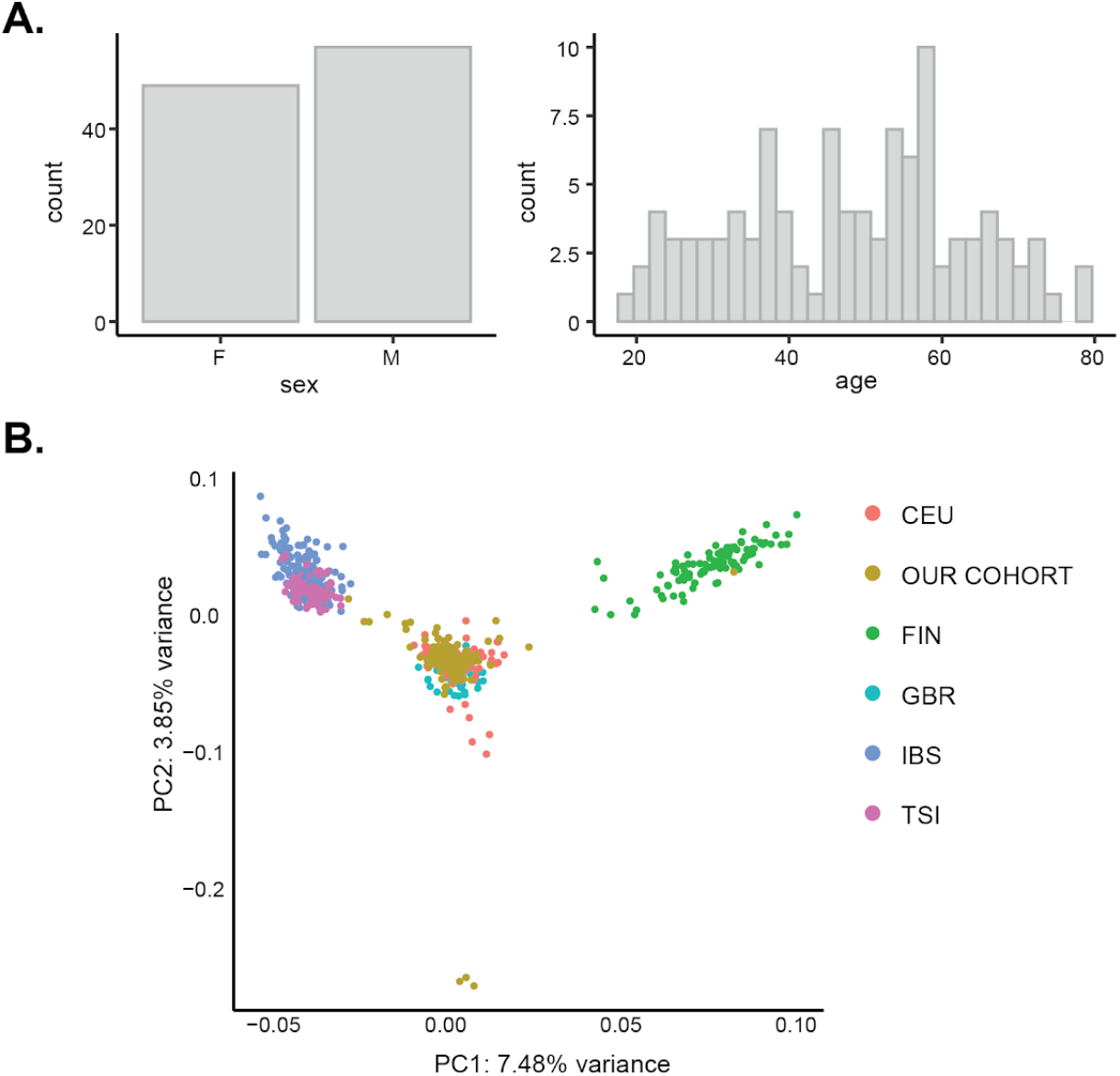
Demographic characteristics of the cohort. **A)** Distriution of sex and age for all individuals. **B)** Genetic ancestry was inferred for each individual in the cohort by comparing their genotypes (common genetic variants obtained from genotyping and imputation) with a panel of five well-defined European ancestries in PCA space. The populations in this panel were derived from the 1000G project (**Methods**) and corresponded to Utah residents with Northern and Western European ancestry (CEU), Finnish in Finland (FIN), British in England and Scortland (GBR), Iberian Population in Spain (IBS), and Toscani in Italy (TSI).

**Supplementary Figure 2.**
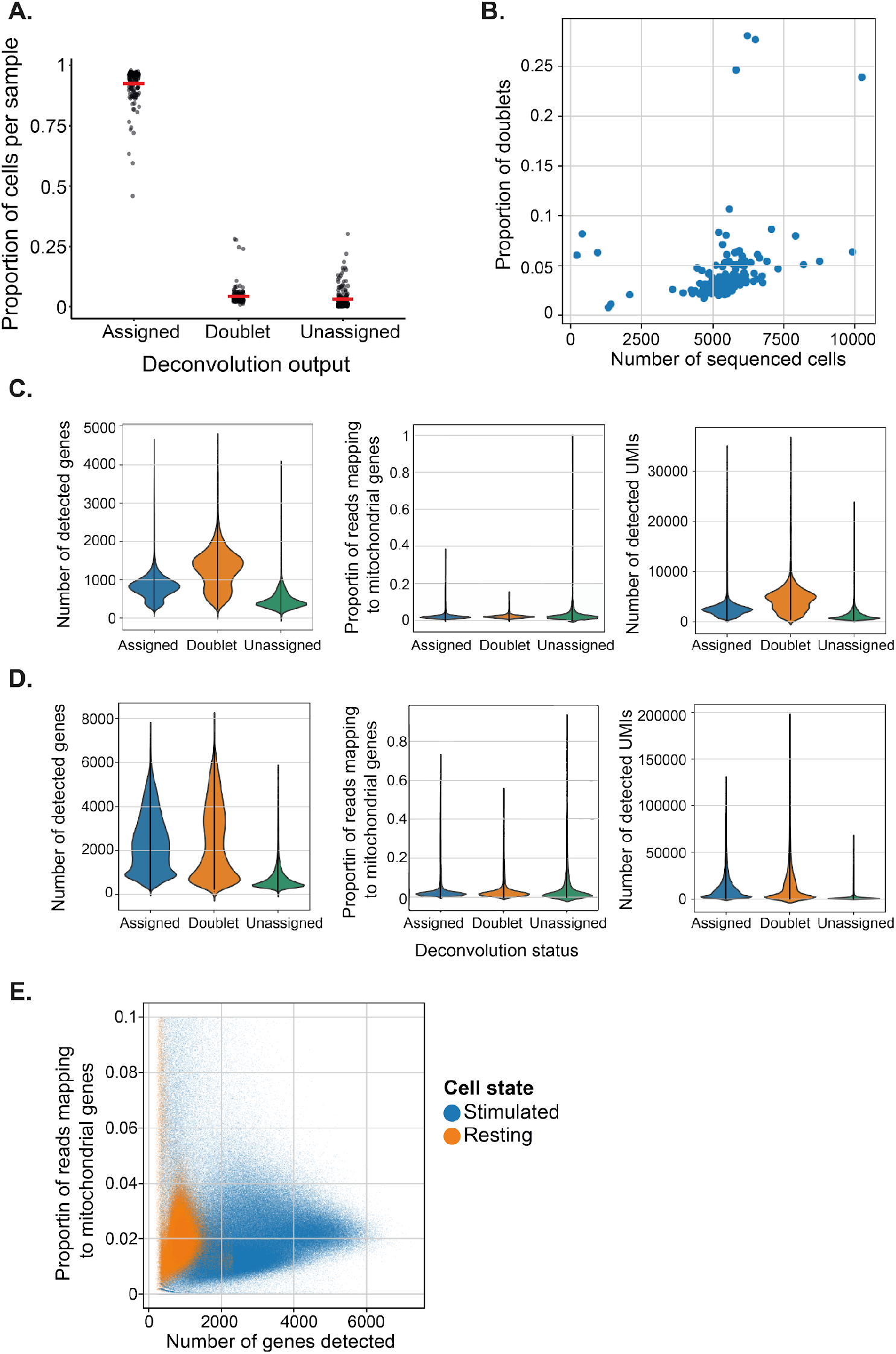
Quality metrics of the scRNA-seq data set. **A)** Proportion of cells per sample which are confidently assigned to one individual (assigned), contain RNA from more than one individual (doublets) or are unassigned (unassigned). **B)** Dot plot showing the relationship between number of cells per inlet and number of doublets. **C-D)** Violin plots showing the number of detected genes (left), proportion of reads mapping to mitochondrial genes (central) and number of detected UMIs (right) in assigned cells, doublets and unsigned cells. These metrics are shown separately for resting (**C**) and stimulated (**D**) samples. **E)** Dot plot showing the number of detected genes and proportion of reads mapping to mitochondrial genes in each cell. Colours indicate resting (orange) and stimulated (blue) cells.

**Supplementary Figure 3.**
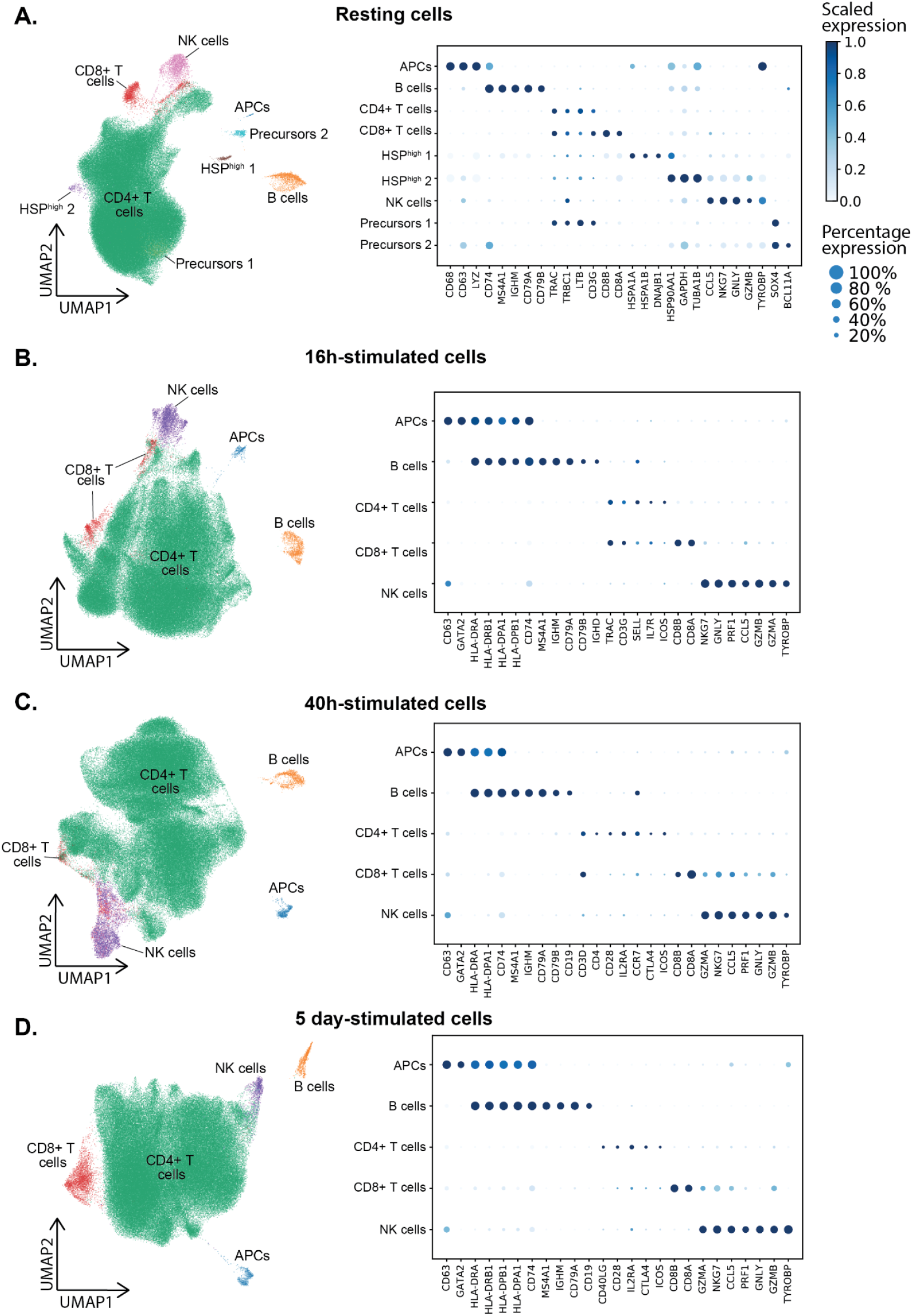
Identification and removal of cellular contaminations in the scRNA-seq data set. Separate UMAP embeddings for cells obtained at four time points. Colours represent cell populations derived from unsupervised clustering. Green cells are CD4+ T cells. Other colours represent contaminations that were removed for subsequent analyses. **A)** Resting cells. **B)** 16h-stimulated cells. **C)** 40h-stimulated cells. **D)** 5 day-stimulated cells.

**Supplementary Figure 4.**
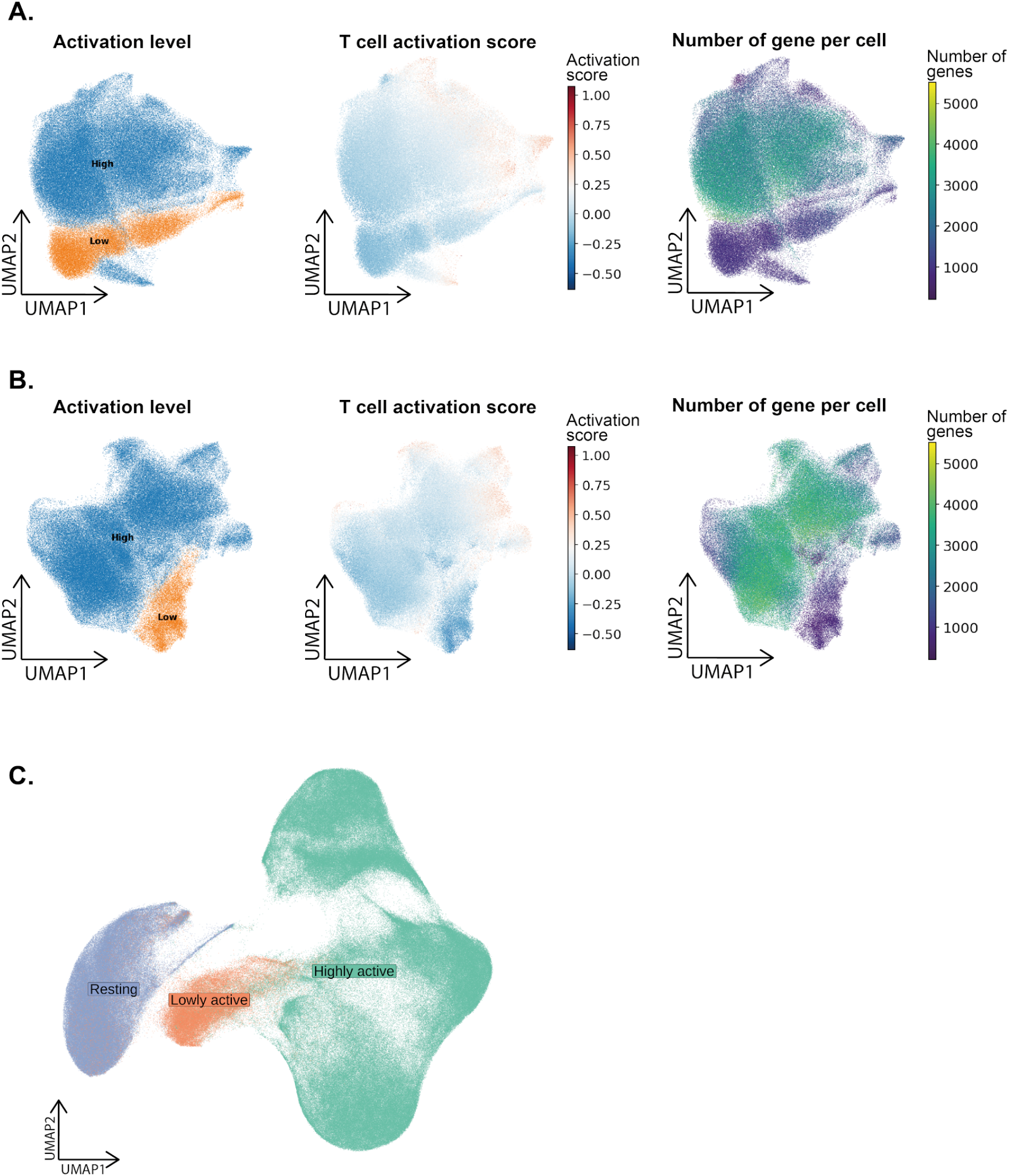
Identification of a population of lowly active CD4+ T cells. A-B) UMAP embedding of all CD4+ T cells sampled at 16h (**A**) and 40h (**B**) after activation. Each dot is a cell, with colours indicating either the lowly active population (orange; left panel), the scaled average expression of a set of T cell activation markers from bulk RNA-seq (central panel), or the number of genes detected per cell (right panel). **C)** Location of the lowly active population of cells in a UMAP embedding containing CD4+ T cells from all time points in the study.

**Supplementary Figure 5.**
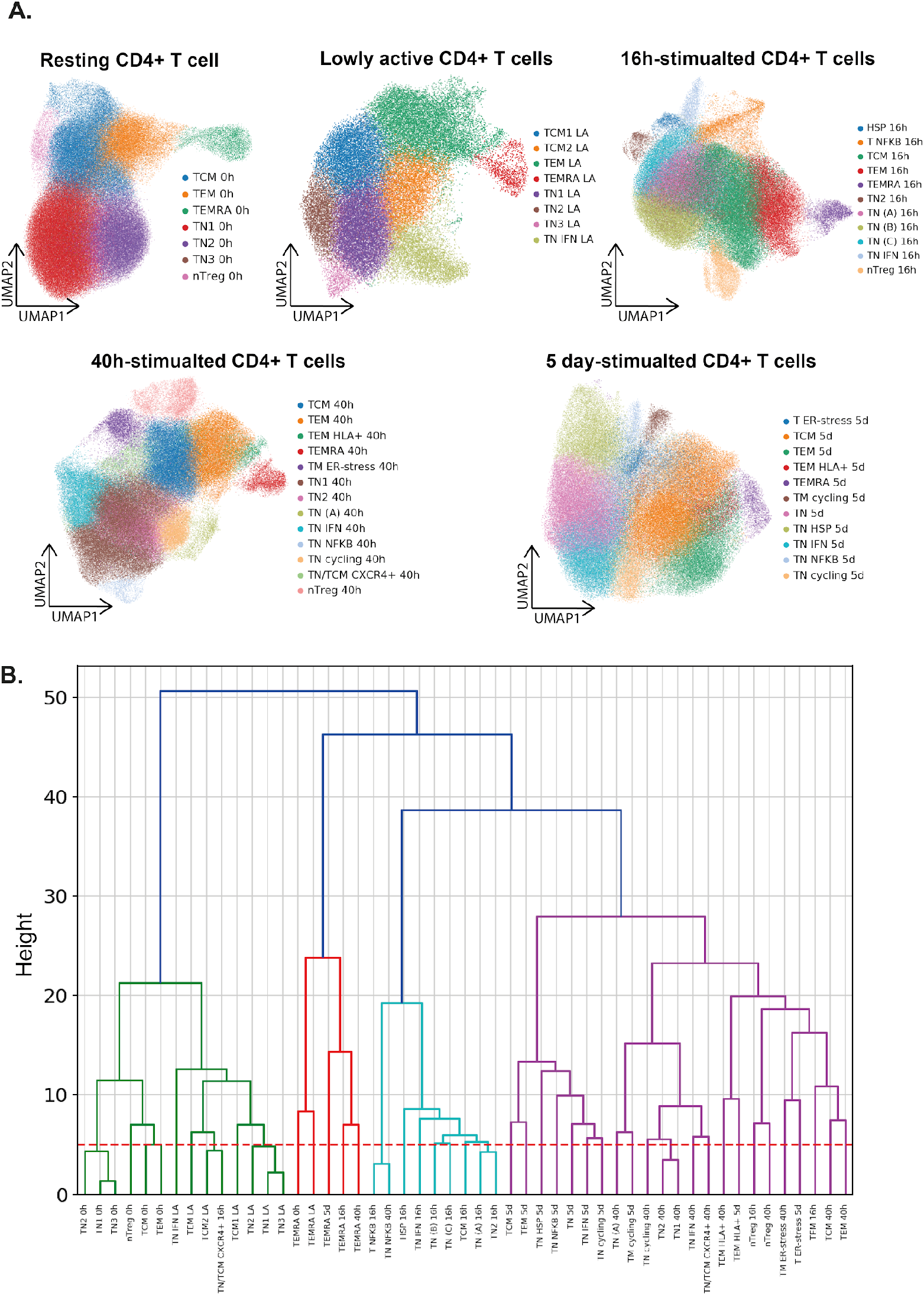
Cluster merging strategy. **A)** Unsupervised clustering was applied independently to the five cell groups of cells identified in the study (resting, lowly active, 16 hours, 40 hours, and five days). Colours represent cell populations derived from unsupervised clustering. This resulted in 51 cell clusters. **B)** The similarity of clusters was assessed by performing PCA and estimating the Euclidean distance between pairs of clusters based on the first 100 principal components. Below the dotted red line are clusters with high levels of similarity that were merged together This resulted in 38 distinct groups of cells.

**Supplementary Figure 6.**
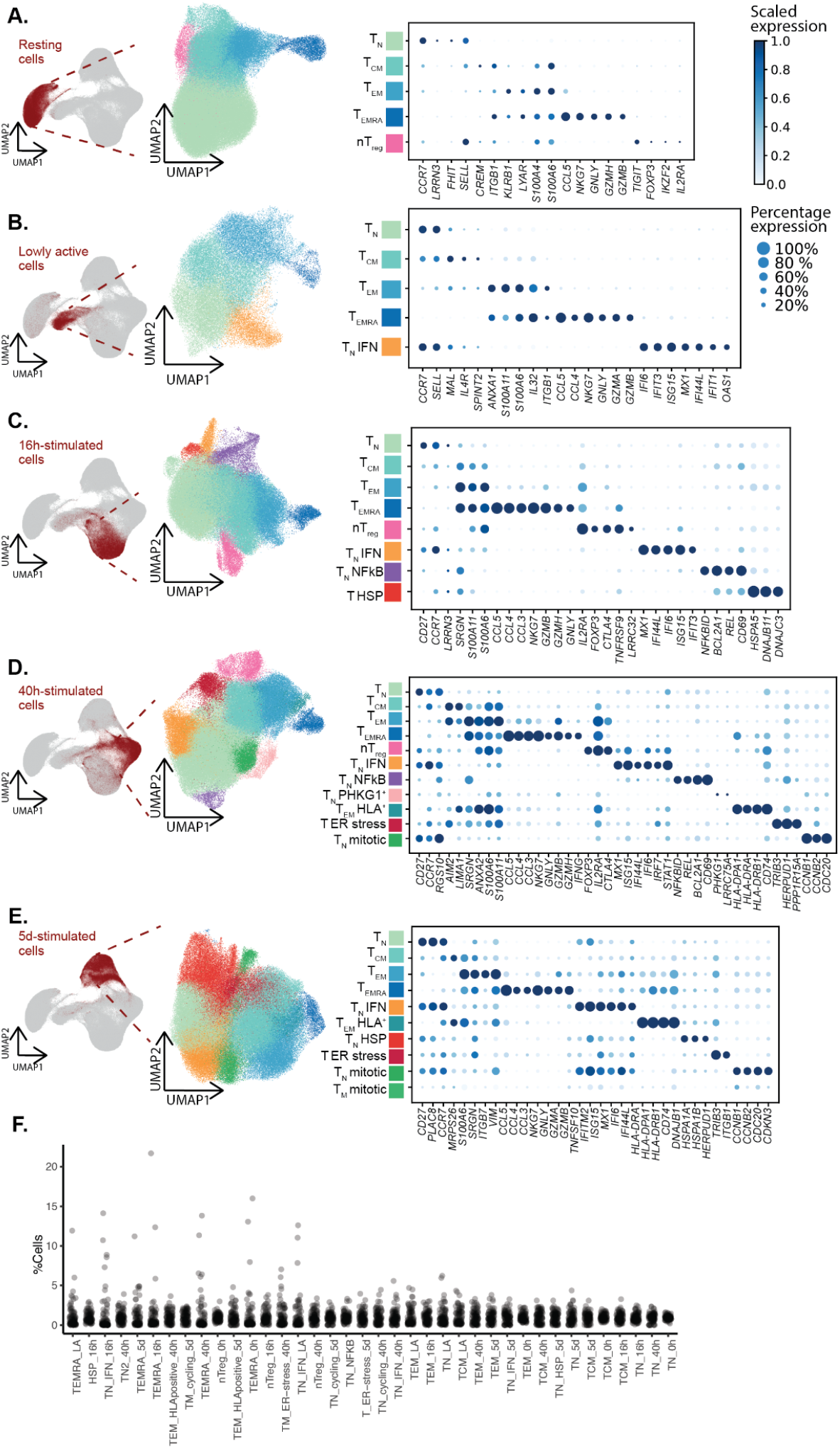
CD4+ T cell subpopulation markers. **A)** Cell subpopulations and marker genes detected independently for CD4+ T cells belonging to each of five broad cell groups: resting (**A**), lowly active (**B**), 16h-stimulated (**C**), 40h-stimulated (**D**), and 5 day-stimulated (**E**) cells. UMAP embeddings (left panels) show each cell as one dot, with colours indicating cell subpopulations derived from unsupervised clustering and cluster merging (38 subpopulations in total). Equivalent subpopulations detected at more than one time point are indicated in the same colour. Dot plots (right panels) show the top marker genes identified for each subpopulation. Shades of blue indicate the scaled mean expression of a gene in each subpopulation, while dot sizes correspond to the proportion of cells in the subpopulation that express the gene. **F)** Dot plot showing the percentage of cells belonging to an individual within each cluster. The X-axis is arranged by cluster size (the smallest to the largest).

**Supplementary Figure 7.**
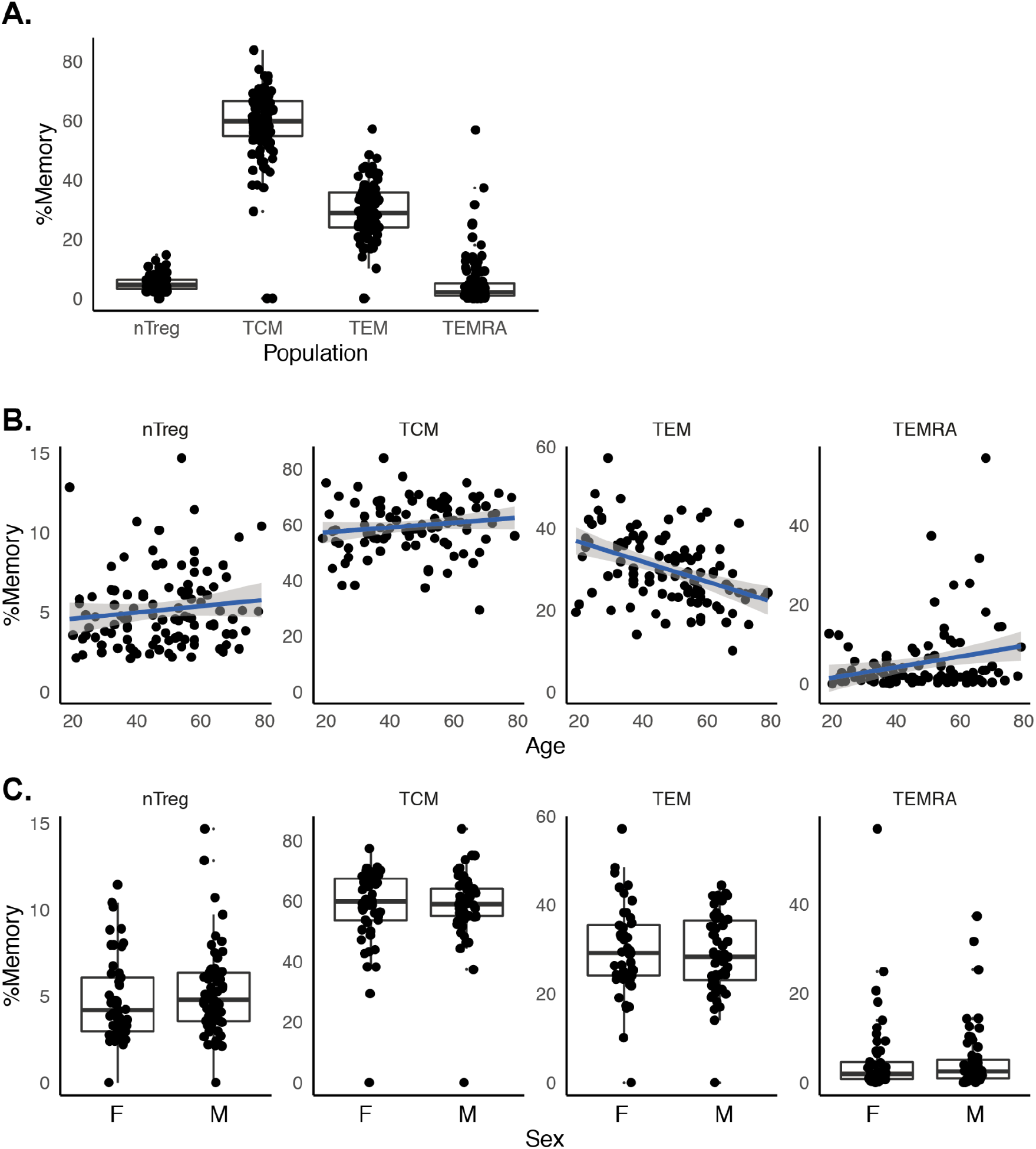
Memory T cell distribution. **A)** Percentage of memory T cell subpopulations. **B)** Relationship between age and percentage of nTregs, T_CM_, T_EM_ and T_EMRA_. **C)** Percentage of memory T cell subpopulations stratified by sex. Each dot represents a measurement obtained from a separate individual.

**Supplementary Figure 8.**
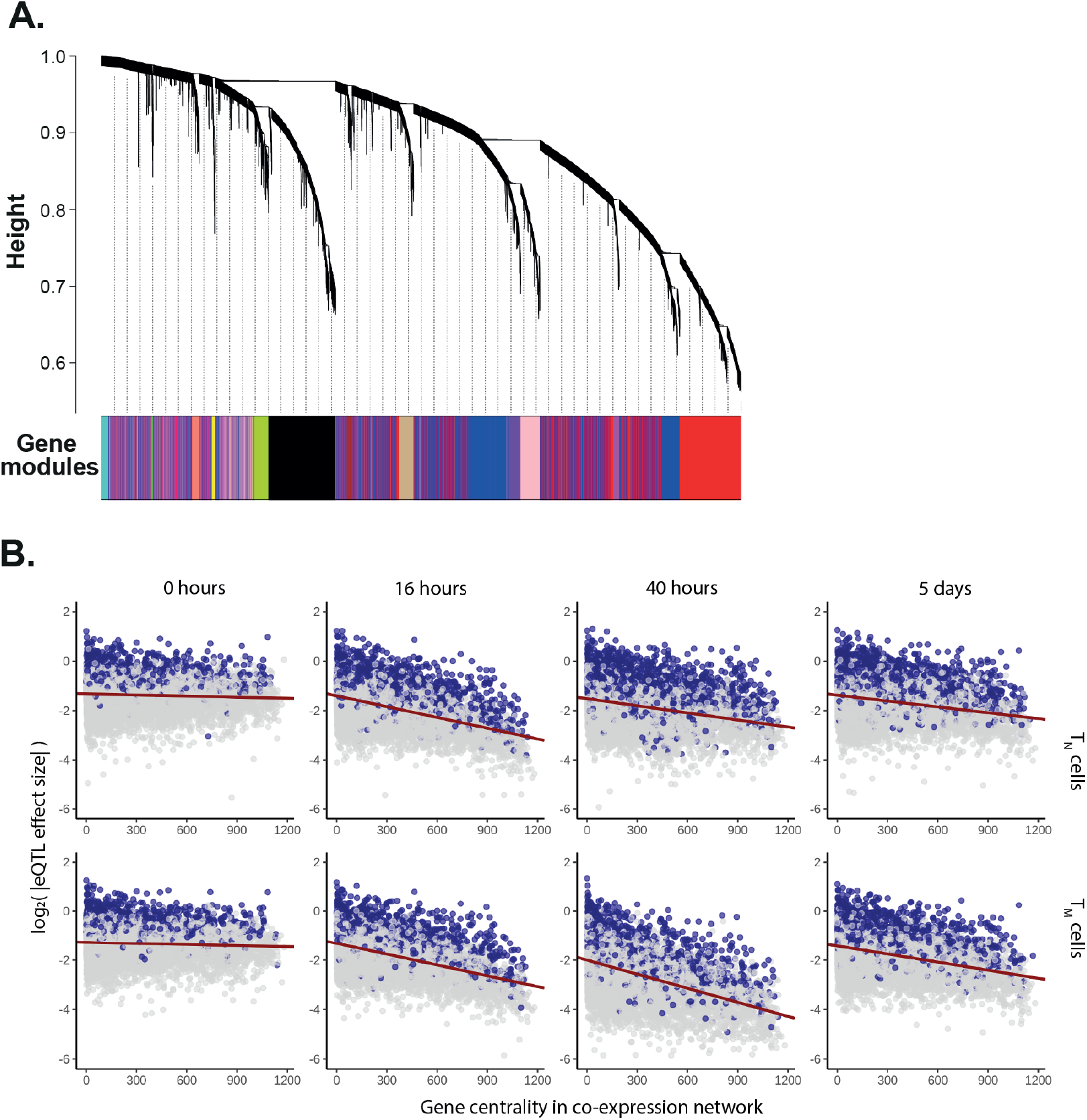
Identification of gene expression patterns active throughout CD4+ T cell activation. **A)** Genes were ordered into a co-expression matrix using WGCNA (Methods). This dendrogram shows genes arranged by similarity. Modules of genes with correlated expression are indicated by blocks of colour at the bottom. **B)** Relationship between a gene’s connectivity (as inferred from co-expression network analysis) and the effect size of its lead eQTL signal. All eQTL effect sizes were log-transformed. Blue dots represent significant eGenes, while gray dots represent genes which do not pass the eQTL multiple test correction.

**Supplementary Figure 9.**
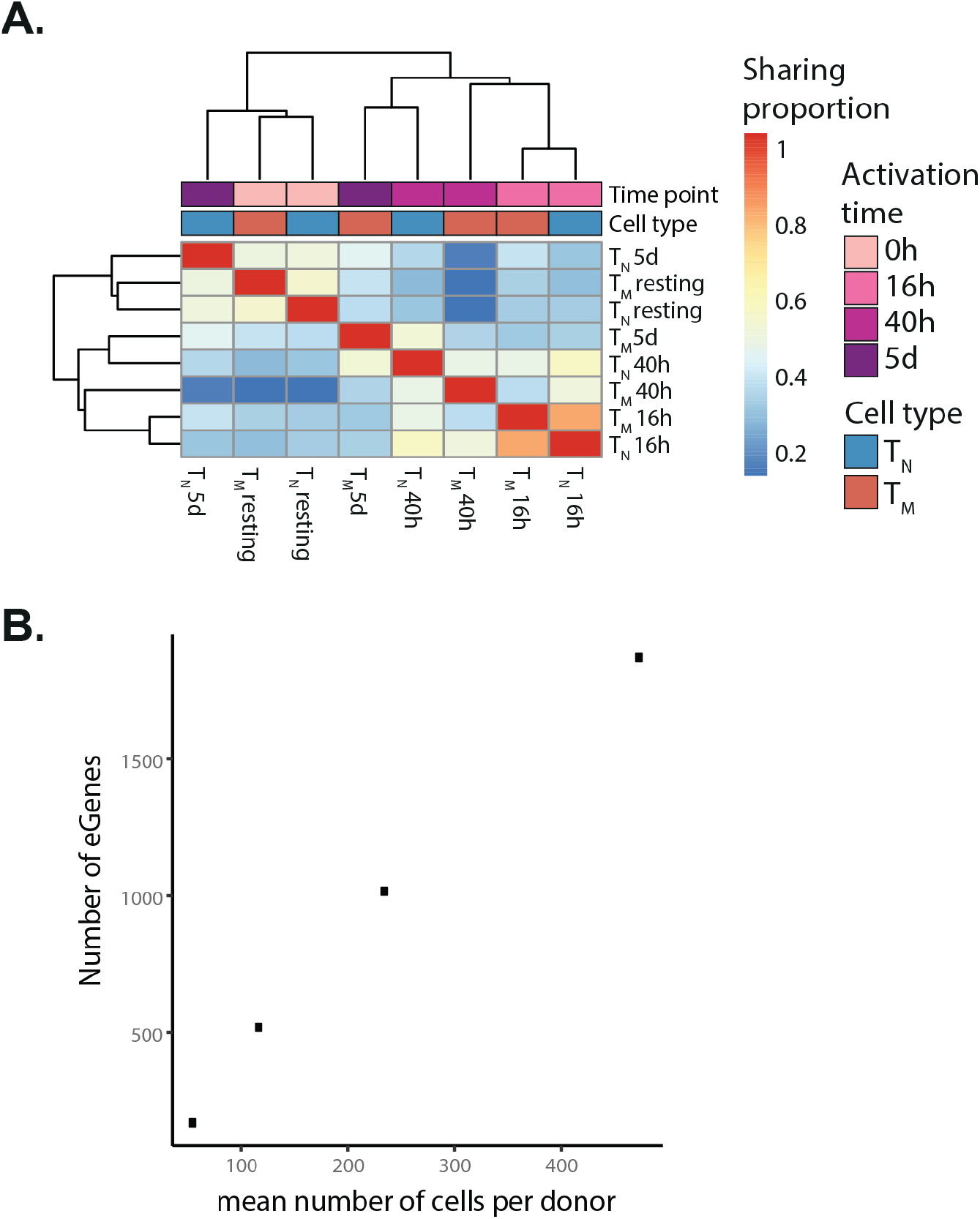
eQTLs sharing. **A)** eQTL sharing was assessed by comparing the effect sizes and directions of eQTLs across cell types (naive and memory cells) and time points using mashR (**Methods**). This heatmap indicates the proportion of eQTLs shared by sign and magnitude between cell type and time point combinations. **B)** The relationship between mean number of cells per donor and power to detect eGenes. T_CM_ cluster was subsampled three times.

**Supplementary Figure 10.**
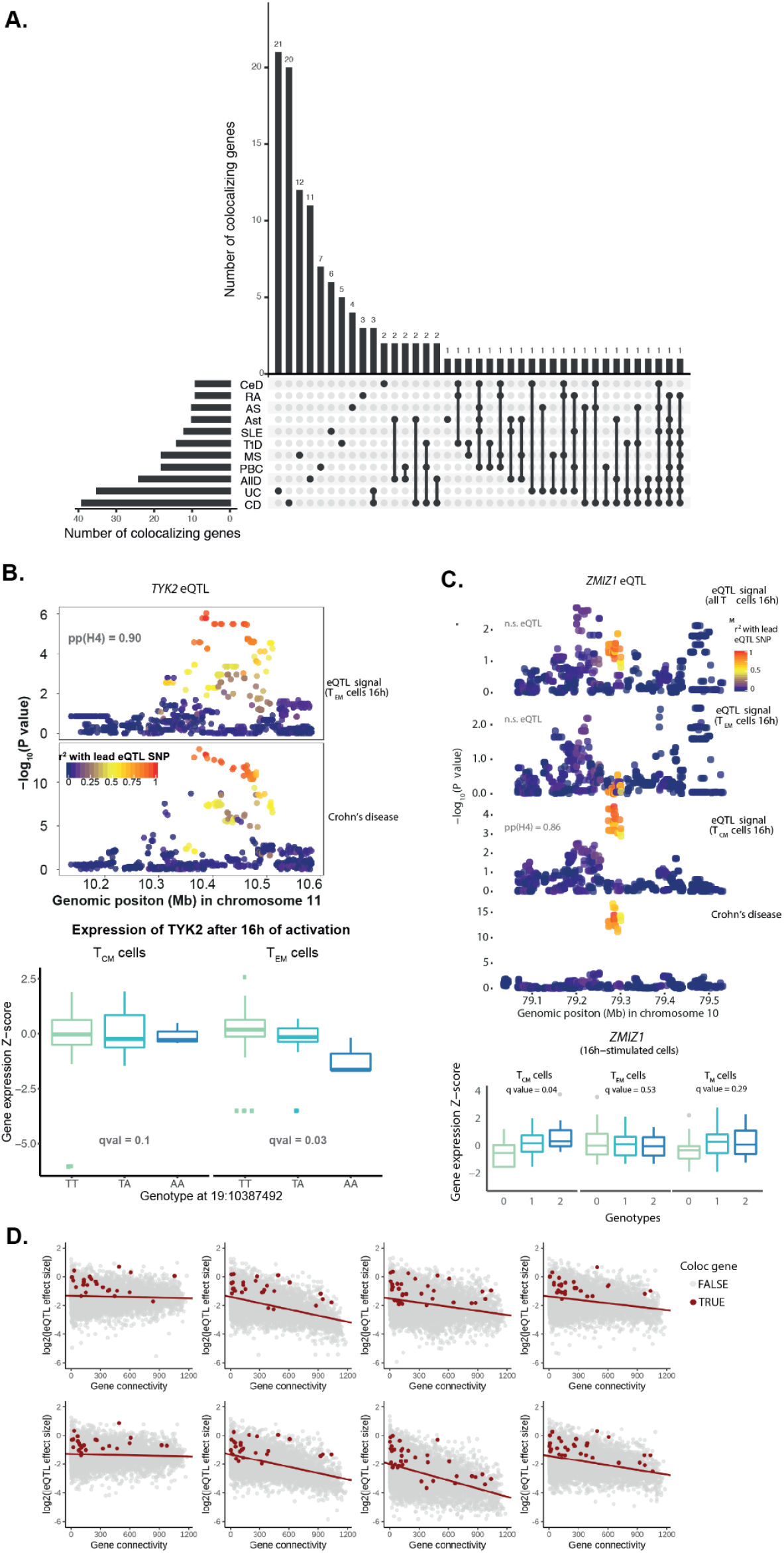
eGenes colocalizing at immune disease loci. **A)** Number of colocalizing genes shared between different immune traits. Bar plots indicate the number of genes in each set. **B and C)** Locus plots for a colocalization between a *TYK2* **(B)** and *ZMIZ1* **(C)** and a GWAS signal for Crohn’s disease. Each dot corresponds to a genetic variant, with colours indicating their LD with the lead eQTL variant. Box plots indicate the expression level of *TYK2* **(B)** and *ZMIZ1* **(C)**, stratified by genotype. Each dot corresponds to a measurement from a different individual. **D)** Relationship between a gene’s connectivity (as inferred from co-expression network analysis) and the effect size of its lead eQTL signal. Red dots represent colocalizing genes and gray dots represent all other genes.

